# Consensus Machine Learning for Gene Target Selection in Pediatric AML Risk

**DOI:** 10.1101/632166

**Authors:** Jenny Smith, Sean K. Maden, David Lee, Ronald Buie, Vikas Peddu, Ryan Shean, Ben Busby

## Abstract

Acute myeloid leukemia (AML) is a cancer of hematopoietic systems that poses high population burden, especially among pediatric populations. AML presents with high molecular heterogeneity, complicating patient risk stratification and treatment planning. While molecular and cytogenetic subtypes of AML are well described, significance of subtype-specific gene expression patterns is poorly understood and effective modeling of these patterns with individual algorithms is challenging. Using a novel consensus machine learning approach, we analyzed public RNA-seq and clinical data from pediatric AML patients (N = 137 patients) enrolled in the TARGET project.

We used a binary risk classifier (Low vs. Not-Low Risk) to study risk-specific expression patterns in pediatric AML. We applied the following workflow to identify important gene targets from RNA-seq data: (1) Reduce data dimensionality by identification of differentially expressed genes for AML risk (N = 1984 loci); (2) Optimize algorithm hyperparameters for each of 4 algorithm types (lasso, XGBoost, random forest, and SVM); (3) Study ablation test results for penalized methods (lasso and XGBoost); (4) Bootstrap Boruta permutations with a novel consensus importance metric.

We observed recurrently selected features across hyperparameter optimizations, ablation tests, and Boruta permutation bootstrap iterations, including HOXA9 and putative cofactors including MEIS1. Consensus feature selection from Boruta bootstraps identified a larger gene set than single penalized algorithm runs (lasso or XGBoost), while also including correlated and predictive genes from ablation tests.

We present a consensus machine learning approach to identify gene targets of likely importance for pediatric AML risk. The approach identified a moderately sized set of recurrent important genes from across 4 algorithm types, including genes identified across ablation tests with penalized algorithms (HOXA9 and MEIS1). Our approach mitigates exclusion biases of penalized algorithms (lasso and XGBoost) while obviating arbitrary importance cutoffs for other types (SVM and random forest). This approach is readily generalizable for research of other heterogeneous diseases, single-assay experiments, and high-dimensional data. Resources and code to recreate our findings are available online.

## Introduction

Acute leukemia is the most prevalent childhood cancer, accounting for 30% of childhood cancers overall [1, 3]. Major subtypes of pediatric acute leukemia include acute myeloid leukemia (AML) and acute lymphoblastic leukemia (ALL), accounting for 15% and 85% of these leukemia cases, respectively [1]. Despite improving survival rates, pediatric AML remains deadlier than ALL [1]. AML is a heterogeneous cancer of the blood and bone marrow myeloid stem cells that presents with numerous molecular subtypes actionable for stratification and treatment. These subtypes are often based on cytogenetics, molecular data, and other characteristics [2, 4]. By contrast to adult AML, pediatric AML is characterized by rare somatic mutations, absence of common adult AML mutations, and relatively frequent structural variants [4]. These findings indicate the importance of age-based targeted therapies for AML treatment, and the potential for molecular assays to further our understanding of how gene expression relates to pediatric AML risk, prognosis, and treatment. We utilized RNA-seq expression data to better understand its relation to pediatric AML risk, which remains poorly understood.

Interest in identification of biomarker and gene target sets of cancer risk using RNA-seq data has endured for over a decade [10]. For statistical rigor and clinical utility, reduction of high-dimensional, whole-genome expression sets of tens of thousands of genes is vital. Differential gene expression (DEG) analysis is typically used to achieve dimensionality reduction by selecting loci with maximal expression contrast between sample groups. This is typically followed by fitting and optimization of models to these reduced sets of DEGs, further narrowing focus to loci showing the greatest contrast and most predictive qualities between sample sets. For the present work, we consider this cumulative process of dimensionality reduction, model fitting, and optimization as a problem of gene feature selection.

Selection of important genes from expression data remains challenging for biomedical research, partly because the commonly applied cross-sectional case/control study design confounds results interpretability. Further, underlying biological dynamics can be nuanced and complex in disease processes, especially for molecularly heterogeneous cancers like AML. These problems can be tractable with modern machine learning approaches, which include the recently developed eXtreme Gradient Boosting (XGBoost) algorithm and Boruta permutation method [7, 12]. With computational advances, these and other methods are more robust, efficient, and accessible to quantitative researchers than ever before. However, these improvements don’t address the need to reconcile disparate findings from applying multiple distinct algorithm types to biomedical data. For this task, it is useful to devise a formalized consensus approach that leverages feature importance metrics across algorithms to arrive at a consensus important feature set. Far from straightforward, development and formalization of consensus feature selection methods with machine learning presents its own challenges. Researchers must reconcile results interpretability, model performance variations, and selection of important features across disparate algorithms and their respective assumptions, strengths, and weaknesses. Further, vital properties of consensus feature selection methods, especially best practices for their use, have yet to be established for biomedical research. Nevertheless, development of such methods is warranted and could become a boon for biomedical research.

The present work is a starting point for addressing aforementioned obstacles for identifying consensus important gene features that help elucidate how gene expression differences relate to pediatric AML risk. We used clinical and RNA-seq data from pediatric AML samples (N = 137 patients) provided by the TARGET consortium. We focused on achieving consensus from 4 distinct algorithms, including lasso, random forest, support vector machines (SVM), and XGBoost [11, 13, 14]. These represent a variety of algorithm types, each with distinct assumptions, strengths, and weaknesses. Random forest and SVM do not natively differentiate important from non-important features, necessitating an importance or weight cutoff be set to identify the most important features. By contrast, lasso and XGBoost perform penalized regression and ensemble learning, respectively, which returns greatly restricted feature subsets, though at the cost of feature exclusion bias (see Results). We addressed these issues by bootstrapping Boruta permutations with a novel consensus importance metric based on relative feature importance rank across these 4 algorithms.

## Materials and Methods

### Pediatric AML Dataset

We accessed TARGET pediatric cancer assay and clinical data from the Genomic Data Commons (GDC, website) on February 4th, 2018. The TARGET pediatric AML cohort consists of samples from 156 patients, with tissues including primary peripheral blood (N = 26), recurrent bone marrow samples (N= 40), primary bone marrow (N = 119), and recurrent peripheral blood (N = 2). For the following analyses, we combined primary blood and bone tissues from 145 patients, retaining one sample per patient.

### Gene Expression Data

RNA-seq data is from pediatric AML patients (N = 137 samples) with clinical and assay data from pediatric cancer patients from the Therapeutically Applicable Research To Generate Effective Treatments (TARGET) initiative, a collaboration between the National Cancer Institute (NCI) and Children’s Oncology Group (COG) clinical trials (website). We obtained RNA-seq expression data as raw gene counts, produced using the Illumina Hi-Seq platform from Genomic Data Commons repository (https://gdc.cancer.gov/). In brief, raw reads were aligned to GRCh38 using STAR aligned in 2-pass mode and gene counts were produced using the HTSeq-counts analysis workflow with Gencode v22 annotations. Full details of the data processing pipeline can be found at the GDC (‘https://docs.gdc.cancer.gov/Data/’). The GDC file manifest are included in (Supplemental Table 4). Gene counts were then normalized using trimmed mean of M (TMM) values method and converted to log2 counts per million (CPM, [18]).

### Pediatric AML Clinical Risk and Binary Risk Classifier

We defined a binarized version of the clinical risk group classifier (low vs. standard or high): Risk group classifications are defined based on patient cytogenetics and mutations, and which pertains broadly to patient outlook in terms of risk of relapse, recurrence, and/or disease progression.

We focused on the “Risk Group” variable from the patient clinical data table. This variable is an aggregate pertaining to a combination of risk of recurrence, progression, and relapse ([1]). Patients were categorized as either low or not-low (e.g. standard or high) risk, and this categorization, called binarized risk group (BRG), was used in the machine learning investigation. Patients missing data for risk group were excluded from the analysis. BRG sample groups were approximately balanced according to important demographic variables, including age at first diagnosis and gender (Table 1).

### Differentially Expressed Genes (DEGs)

To reduce noise and false positive rate, we opted to exclude genes with low expression levels and which demonstrated significant differential expression in a contrast between the binarized risk groups in the training data subset using the voom function from the limma Bioconductor package ([11]). With this pre-filter, we identified N = 1,998 (9.33% retained) differentially expressed genes (DEGs) showing substantial mean differences between risk groups (absolute log2 fold-change ¿= 1, adj. p-value ¡ 0.05).

### Machine Learning Algorithms and Hyperparameter Optimizations

We trained and tested gene expression-basd models for predicting BRG using a variety of algorithms, including two types of ensemble approaches (random forest and XGBoost), a kernel-based classifier (Support Vector Machines or SVM), and penalized regression (lasso). These algorithms quantify feature importance in the following ways: 1. Lasso assigns beta-value coefficient (positive, negative, or null/0) for use in penalized regression; 2. SVM assigns a feature weight (positive or negative) for inclusion in kernel-based estimator; 3. XGBoost assigns importance (positive or null/0) from gain across splits; 4. Random forest assigns importance using mean decrease in Gini index (positive value).

With each algorithm type, we fitted models by varying algorithm hyperparameters (Table 1, Figure 2, and Results). For Random Forest, we varied the number of trees (ntrees) from 2,000 to 10,000. For XGBoost, we varied training depth and repetitions. For SVM, we varied the kernel type to be linear or radial, and the weight filter to be none or 50%. For lasso, we varied the alpha value to be from 0.8-1.2 (Table 1 column 3). These runs informed hyperparameters used in each of the 4 algorithms with bootstraps of Boruta permutations (Supplemental Material, Figure 4).

### Permutations of Sample Label Switching

To test accuracy of sample labels and quantify possible miss-classification, we performed permutation tests with risk label reassignment. For each algorithm, the training dataset class labels were randomly permuted (switched) 5000 times, such that each patient in the training set was randomly assigned to, the class label switching allows one to infer that the feature contribution for correct classification is not likely due to chance.

### Ablation Tests

To characterize predictive gene sets and networks, we performed ablation tests with penalized algorithms (lasso and XGBoost). In each ablation iteration, we excluded selected gene features from all prior iterations before re-fitting and assessing fitted models with remaining DEGs. We repeated this for 15 and 70 iterations for lasso and XGBoost, respectively (Figure 3, Supplemental Figures 1 and 2, Supplemental Materials). We assessed the expression correlation (whole sample dataset) between first iteration selected genes and the next successive 2 and 3 iterations for lasso and XGBoost, respectively (Figure 3B and 3C, Supplemental Figures 1 and 2).

### Analysis Code and Data Availability

Analysis was conducted on the publicly available TARGET pediatric AML cohort (Supplemental Table 4 for download manifest). The majority of analysis was conducted using the R programming language with packages from Bioconductor and CRAN repositories ([7, 11-14], Supplemental Methods). Pediatric AML RNA-seq and clinical data were bundled into SummarizedExperiment objects for convenience (Supplemental Materials). Scripts, notebooks, code, and data objects are available online (website).

## Results

### Pediatric AML Risk Group Demographics

This study focused on whether gene expression could be used to predict pediatric AML risk, as defined using the classical cytogenetic and molecular classification scheme [3]. We initially identified TARGET pediatric AML patients with primary blood or bone cancer samples (N = 137 patients) and defined a binarized risk group (BRG) classifier as either low risk or not-low risk, where the latter category combines “standard” and “high” risk patients (Figure 1). Summary statistics indicated binarized risk was approximately balanced for important demographic characteristics, including age at first diagnosis, gender, and bone marrow leukemic blast percentage and peripheral blasts (Supplemental Table 1). However, the “not-low” risk group had a significantly lower median for white blood cell count at diagnosis (29.3 [range: 1.30-519] in not-low versus 53.5 [range:1.60-253] in low-risk, p = 0.032). We randomly divided samples into training (N = 96 samples) and test (N = 49 samples) subsets, at a ratio of 2:1, preserving BRG sample group frequencies in each subset. We used training data to calculate differentially expressed genes (DEGs), and the train and test set classifications to fit and assess fitted models below.

**Figure 1.**
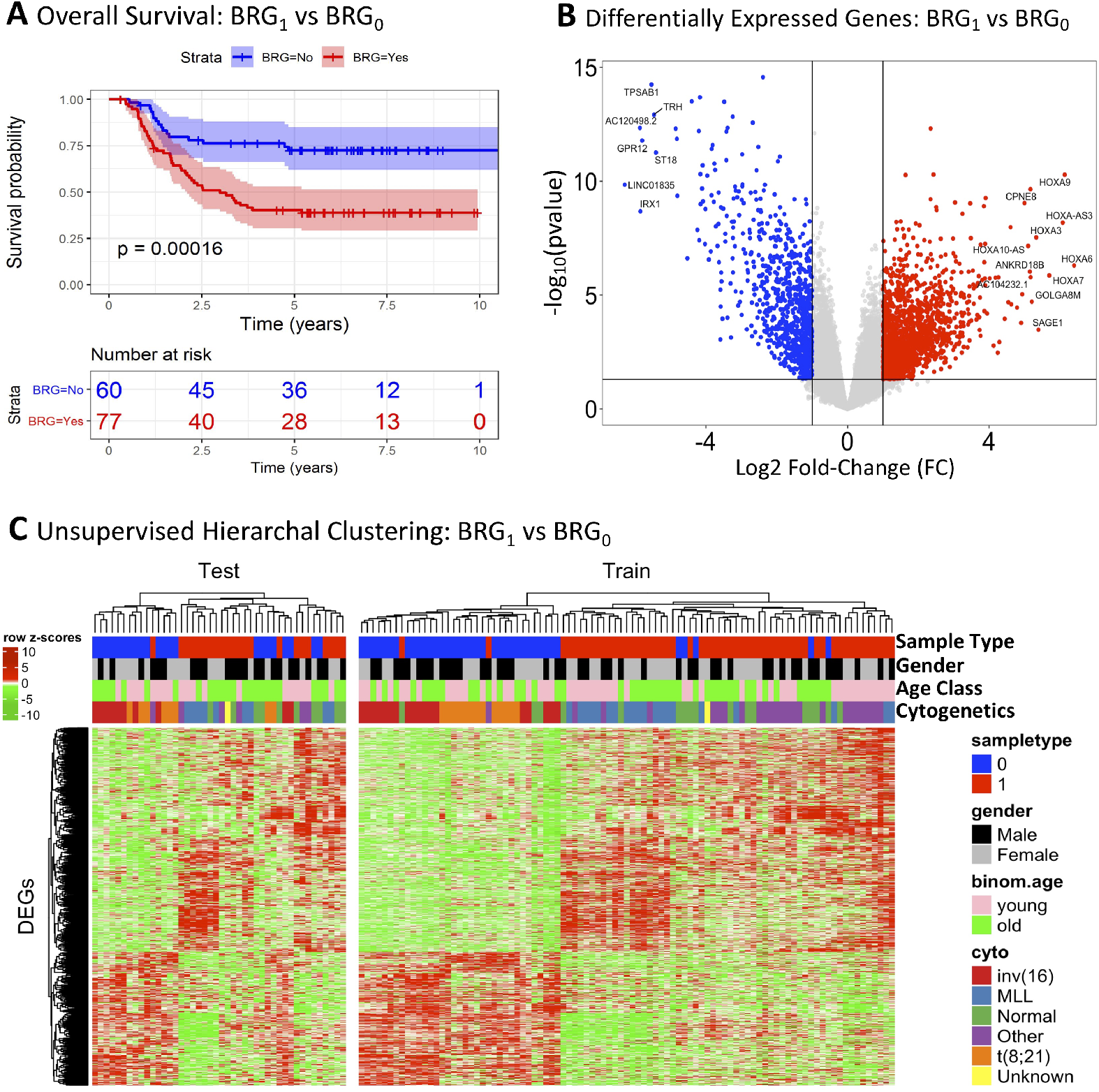
Clinical and demographic information summary for TARGET pediatric AML dataset. A. Survival in Low (binary risk group, BRG = 0), compared to Not-Low (BRG=1) clinical risk group. B Volcano plot of differentially expressed genes (DEGs), x-axis is log2 fold-change, y-axis is −1 times log10 of unadjusted p-value from t-tests, significance threshold (horizontal line) set at ¡0.01 p-adjusted and (vertical lines) —log2FC—¿1. C Heatmap of DEG expression (Z-score of normalized expression) with sample-wise clinical annotations (“cto” is primary ctogenetic subtype).

### Dimensionality Reduction with Differentially Expressed Genes (DEGs)

We pre-filtered the RNA-seq gene expression dataset to limit the number of features included in the initial model training. Using the training dataset, gene expression for standard or high risk patient (not-low risk, BRG = 1, N = 55 samples) were contrasted to patients at low risk (low risk group, BRG = 0, N = 38 samples) using differential expression analysis. From approximately 60,000 genes assayed, we identified 1,984 differentially expressed between risk groups (—log2FC— <1, p-adj. >0.05, Figure 1B and 1C, Supplemental Table 2, Methods). This increased the mean of normalized expression differences from 0.50 to 1.71 (median increase from 0.32 to 1.51). Mean of variance differences also increased from 0.76 to 2.19 (median increase from 0.31 to 2.19).

### Algorithm Hyperparameter Optimization

We performed hyperparameter optimization with four distinct algorithm types (lasso, random forest, SVM, and XGBoost) to determine optimal values to use in following ablation and consensus tests (Figure 2, Table 1). These algorithms include two ensemble methods (random forest and XGBoost) two penalized methods (XGBoost and lasso) and two unpenalized methods (SVM and random forest). These algorithms quantify feature importance in the following ways: 1. Lasso assigns beta-value coefficient (positive, negative, or null/0) for use in penalized regression; 2. SVM assigns a feature weight (positive or negative) for inclusion in kernel-based estimator; 3. XGBoost assigns importance (positive or null/0) from gain across splits; 4. Random forest assigns importance using mean decrease in Gini index (positive value). For each algorithm, we tested at least 3 distinct hyperparameter value sets (Table 1 column 3), and compared model performances. We observed a variety of model performance fluctuations across optimizations for each algorithm (Supplemental Material). All fitted XGBoost and lasso models showed uniformly high performance. For SVM, radial kernel tests showed worse performance than linear kernel tests. Where there were clear performance benefits, we selected the optimal hyperparameter sets for inclusion in our consensus importance metric (Figure 4, Supplemental Material).

**Figure 2.**
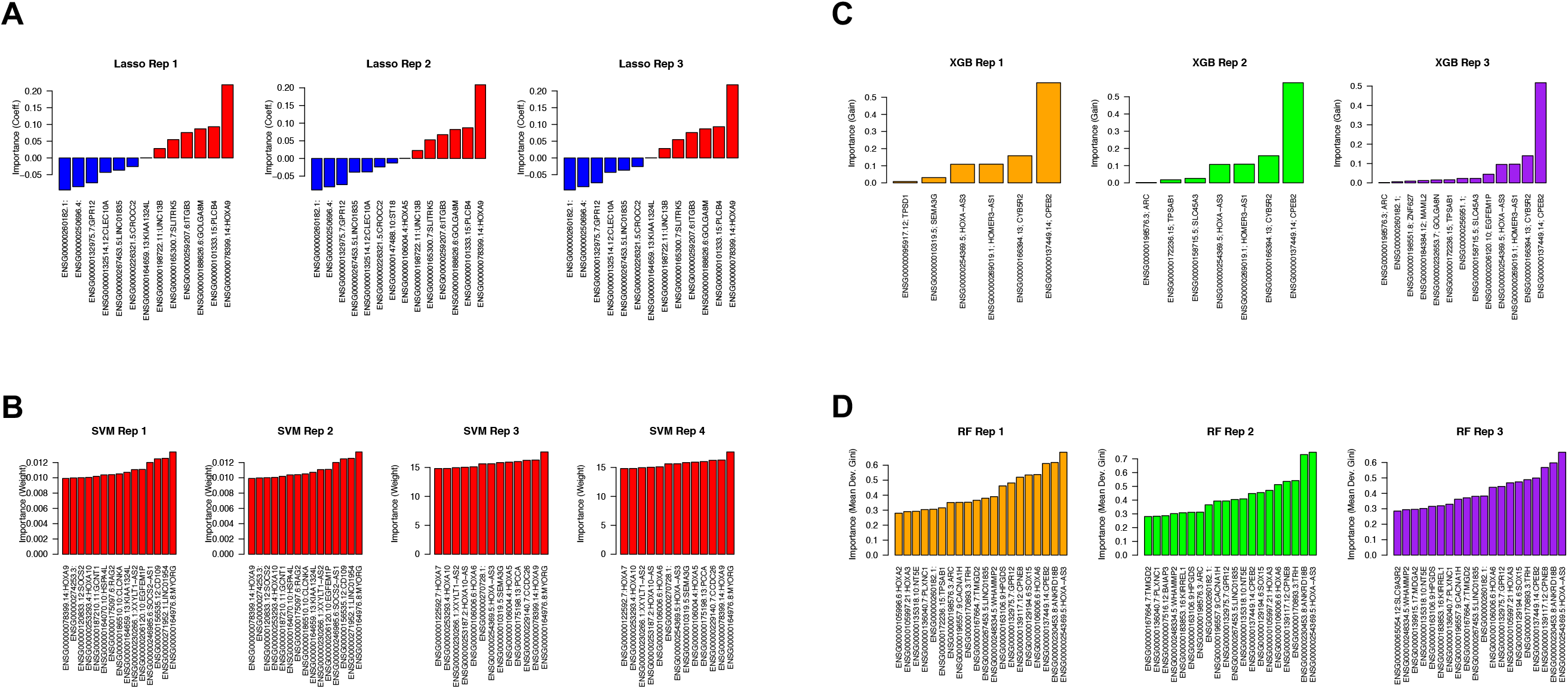
Results of hyperparameter optimizations. A Results of 3 lasso iterations varying alpha from 0. 5-1.2 (shows all genes with not-null coefficients). B SVM 4 iterations, varying linear and radial kernel, and no weight filter versus top 50% weight filter (shows features with top 99th quantile absolute weight). C XGBoost (“XGB”) 3 iterations, varying steps (shows all with not-null importance). D Random forest (“RF”) 3 iterations varying ntrees from 5-15k (shows top 99th quantile importance).

**Figure 3.**
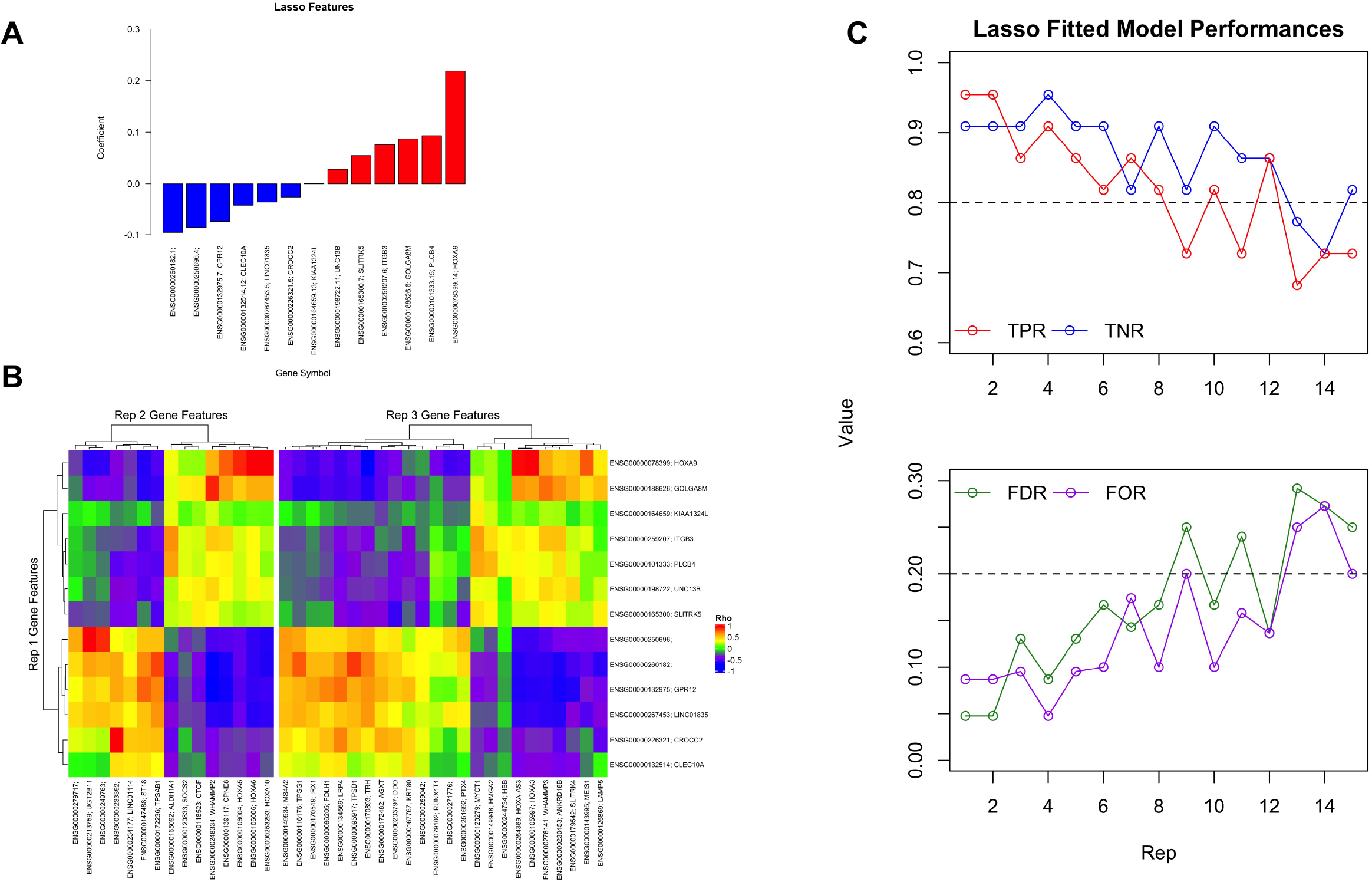
Lasso ablation test results. A. Beta-value coefficients (non-zero) from first lasso iteration. B. Correlation of selected gene feature expression (Spearman Rho, whole dataset) from iterations 2 and 3 with iteration 1 features expression. C. Fitted model performance across 15 ablation iterations, showing (top) true positive (TPR, red) and true negative (TNR, blue) rates, and (bottom) false discovery (FDR, green) and false omission (FOR, purple).

**Figure 4.**
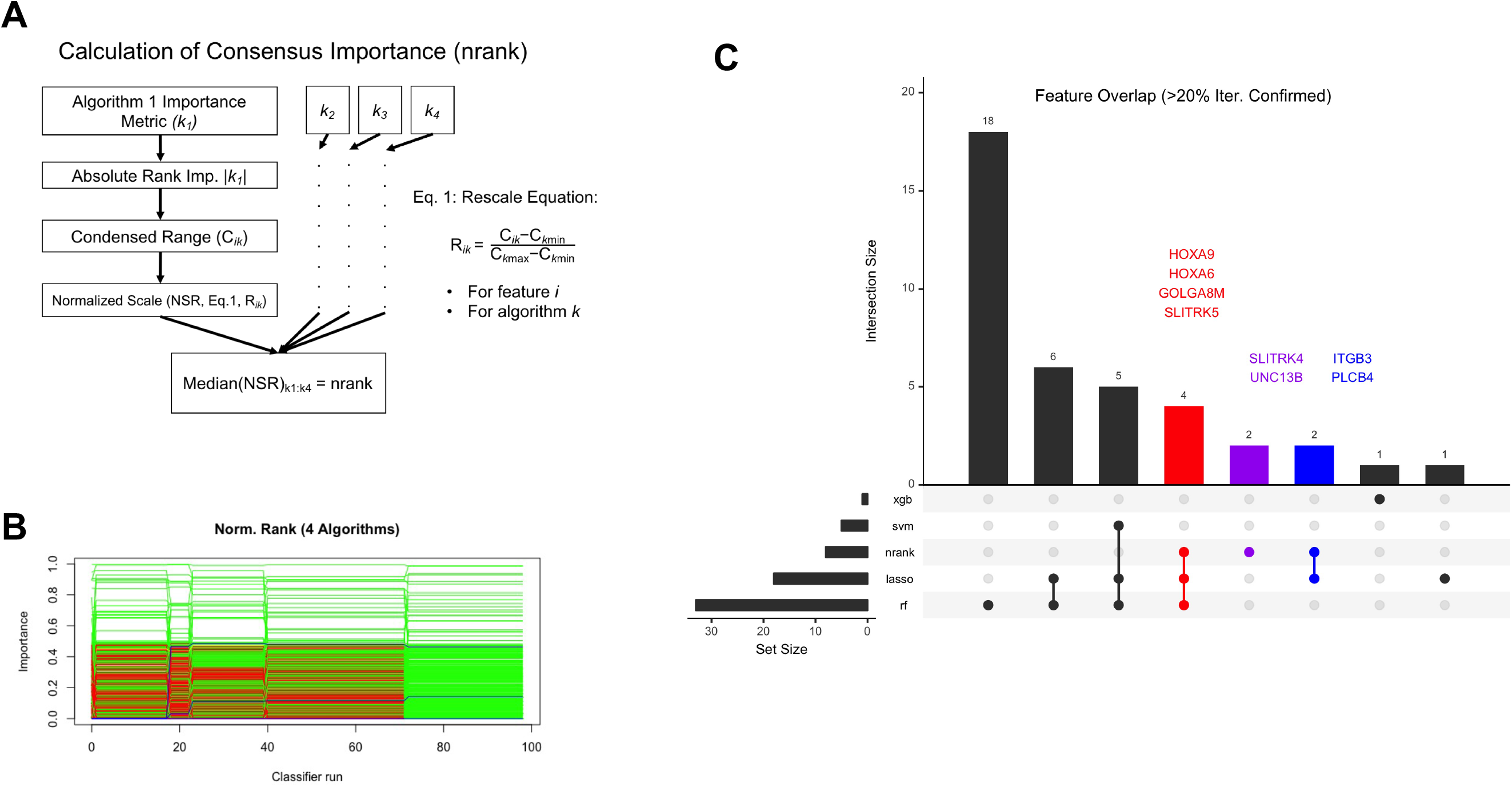
Methods to determine consensus important gene features from Boruta bootstraps (N = 1,000). A Workflow calculating “nrank” consensus importance, or normalized median absolute importance rank, across 4 algorithms (lasso, SVM, random forest, and XGBoost). B Feature (green is confirmed, red is rejected, yellow is tentative) and shadow feature-wise (blue lines) importance (rank, y-axis) across Boruta permutations (x-axis, max = 100). C Upset plot of recurrent confirmed features (present in at least 20% or 200/1,000 bootstraps) across Boruta bootstrap analyses with 5 distinct importance metrics (XGB = XGBoost importance, SVM = SVM importance, Nrank = consensus importance nrank, Lasso = lasso importance, RF = random forest importance). Red genes and data are shared across consensus, lasso, and random forest runs, purple is confirmed genes unique to consensus run, blue is confirmed genes shared only by consensus and lasso runs.

### Ablation Tests and Exclusion Bias with Lasso and XGBoost

Unlike Random Forest and SVM, lasso and XGBoost penalize uninformative and/or correlated features, resulting in 0 or null importance assignment for most genes. Lasso assigns a beta-value coefficient for regression, and XGBoost estimates gain from fractional contributions to splits. Feature omission can reduce data dimensionality and overfitting risk, though this is likely not optimal in biomedical research settings where the objective is to identify a set of gene targets. Exclusion of correlated features can obscure gene sets or pathways of importance, constituting an omission bias. We performed ablation studies using lasso and XGBoost. For each iteration of ablation, we excluded all features selected from prior iterations before refitting lasso or XGboost models, respectively (Figure 3, Supplemental Figures 1 and 2, Supplemental Materials, Methods).

In the absence of an omission bias, we expected consistent decline in fitted model performance with successive ablation iterations. Instead, we observed oscillation between performance recovery and decline across successive ablation iterations, with gradual performance decline across 15 and 70 ablation iterations of lasso and XGBoost, respectively (Figure 3C and Supplemental Figure 1). Interestingly, models fitted in later iterations could recover substantial performance, and this trend was even more exaggerated for XGBoost than lasso ablation iterations. This trend likley reflects signal gain and loss of alternative predictive and related or correlated gene sets and pathways, which are unrepresented in sets from earlier ablation iterations. As iteration increases, gene members of alternate functional sets may be successively selected then exhausted, resulting in initial performance recovery followed by successive performance loss. These findings highlight the importance of carefully evaluating iterations of penalized methods in biomedical research, and the utility of ablation tests.

We observed substantial correlated expression, both positive and negative, across genes selected in the first 3 and 4 ablation iterations for lasso and XGboost, respectively (Figure 3C and Supplemental Figure 2). Correlated expression could result from direct or indirect functional interactions or relatedness. We observed evidence for functional similarity across these selected gene sets, especially shared HOX pathway membership. Surprisingly, HOXA9 was selected in the first iteration of lasso ablation, but not until the fourth iteration of XGBoost ablation (Figure 3A, Supplemental Figure). HOXA9 is known to be co-expressed in multiple pediatric AML subtypes, and its activity can be used to predict patient risk [5, 9] (Figure 3A). We further note substantial positive correlation between HOXA9 and the HOX family gene MEIS1, which was selected in iteration 3 of lasso ablations. MEIS1 expression is linked to hematopoietic stem cell development, and HOXA9-MEIS1 complexes were found to correlate with AML subtype and outcome [8, 17, 19, 20].

### Consensus Important Gene Features from Boruta Permutation Bootstraps

We designed and applied a consensus machine learning algorithm to identify recurrent important gene features. We used a consensus importance metric (”nrank”, Figure 4, Supplemental Figure 3-10, Supplemental Material), which returns a normalized median absolute rank after calculating the algorithm-specific importance metrics from lasso, random forest, XGBoost, and SVM. We then permuted this calculation in the Boruta method for 1,000 bootstraps, with redraw of 2/3rds of pediatric AML samples in each bootstrap, to simulate redraw of the training sample subset ([12], Supplemental Methods). For comparison, we also used single-algorithm importance for 4 algorithms in Boruta permutations across 1,000 bootstraps apiece (Supplemental Material). Across these tests, we evaluated importance calculation histories (Supplemental Figure 4), gene-wise summaries of label assignments (either “rejected”,”tentative”, or “confirmed” in each Boruta bootstrap iteration, Supplemental Figures 5-9), and finally extent of consensus across Boruta bootstrap runs using each respective importance metric (Figure 4C, Supplemental Figure 10).

To understand these results, it is necessary to summarize the Boruta method. In each permutation, this method calculates observed importance for “real” features and importance distribution of “shadow” features. Shadow features are obtained by random reassignment of expression values to samples, which breaks correlation of expression with class (AML risk). Real features are rejected if their importance is sufficiently similar to the shadow feature importance distribution. Remaining features are then retained in following permutations. Ultimately a label of “rejected” (non-important), “tentative” (marginal features), or “confirmed” (high-confidence important features) to each gene feature (Boruta citation). Evaluating real and shadow feature importance across Boruta permutations for a sampling of bootstraps, we observed a range of behavior across the various importance metrics used (Supplemental Figure 4). XGBoost showed no exclusion of rejected features across permutations. By contrast, the remaining methods, including nrank consensus, showed progressive retention of confirmed or tentative features and omission of rejected ones.

We studied recurrent selected genes in each test by setting progressively more stringent cutoffs (e.g. gene was labeled confirmed in >1, >20% or 200/1,000 bootstraps, or >50% or 500/1,000 bootstraps). We observed a range in the total sizes of confirmed feature sets across runs, and total recurrent confirmed feature sets from the consensus nrank run fell in the middle of this range. Interestingly, about 50% of consensus nrank confirmed features overlapped with recurrent confirmed features from random forest, SVM, and lasso runs, though not XGBoost (Figure 4C, Supplemental Figure 10). Certain confirmed genes, including HOXA9 and MEIS1, were present in the final recurrent confirmed gene set (Table 2). These genes were identified in the first 4 ablation iterations with lasso and XGBoost, and their inclusion in the consensus gene set indicates our approach mitigates exclusion bias of independent penalized algorithms.

## Discussion

We present a consensus feature selection strategy, including a novel consensus rank importance metric and implementation with bootstraps of Boruta permutations. This consensus approach can mitigate possible algorithm feature exclusion biases of penalized algorithms (lasso and XGBoost) while obviating the need to set arbitrary importance cutoffs with algorithms not natively performing feature selection (random forest and SVM). Prior studies characterized numerous molecular subtypes in pediatric AML, reflecting heavy utilization of whole genome sequencing, methylation, and other assay types, with less utilization of RNA-seq gene expression data ([4, 6]). We focused primarily on data from RNA-seq, which may be underutilized for characterization of pediatric AML and clinical risk. Our consensus gene feature set validates prior literature and demonstrates how a single-assay approach can be used to characterize clinical risk in a molecularly heterogeneous cancer.

Among consensus important genes for pediatric AML risk, we identified numerous potential therapeutic targets. HOXA9 has been implicated in MLL (KMT2A)-rearranged AMLs and MLL-HOXA9 fusion has been shown to induce leukemogenesis in xenograph and mouse models [16]. Interestingly, SLTRK5 and ITGB3 are both highly and aberrantly expressed on the cell-surface. This suggests these genes may be good potential targets for antibody and CAR-T cell therapies. SLTRK5 has been shown to be aberrantly expressed in nearly 80% of AML and coincides with high/standard risk clinical features, allowing one the potential to improve outcomes for AMLs with poor prognosis [15].

While the present work describes a thorough application of algorithm and machine learning to elucidate expression-based gene targets in AML risk, we must note certain limitations. The TARGET pediatric AML dataset is moderate in size, and further limited when training and test subsets are used, increasing risk of model overfitting. We note that our consensus approach could mitigate the effects of individual model overfitting because its output is a gene list rather than a fitted model.

Our consensus machine learning approach can and should be formalized and fine-tuned for better performance, efficacy, and generalizability. This can be achieved in several possible ways, including iterative recalculation of DEGs, bootstrapping in hyperparameter optimization, and inclusion of alternative consensus importance metrics besides the “nrank” method used here (Results, Supplemental Material). Finally, best practices for consensus feature selection, such as minimal data size or optimal test/train split for a given effect magnitude, have yet to be established for biomedical data. To this end, our results here are promising, and we have provided sufficient notebooks and scripts such that our consensus method can be generalized to research other diseases or datasets.

## Supporting information

Main Tables

Boruta Methods Notebook

## Supporting Information

**S1 Figure**

**Correlation of gene expression (entire dataset) across gene features selected in iteration 1 vs. iterations 2-4 of XGBoost ablation tests (see Results, Supplemental Materials).**

**S2 Figure**

**XGBoost fitted model performances across 70 iterations of ablation. (Top) true positive rate (TPR), and true negative rate (TNR). (Bottom) false discovery rate (FDR) and false omission rate (FOR).**

**S3 Figure**

**Boruta permutations consensus rank metric comparison. Comparison of naive (x-axis) and normalized (y-axis) rank across features, for each of 4 algorithms used (see Figure 4A, Supplemental Materials).**

**S4 Figure**

**Comparison of feature importance across Boruta permutations for 5 bootstrap iterations (selected at random). Each column shows bootstraps for permutations with a different importance metric (either lasso, consensus importance “nrank”, random forest “rf”, SVM “svm”, or XGBoost “xgb”).**

**S5 Figure**

**Feature classification summary across Boruta bootstraps (N = 1,000) with consensus importance (“nrank”).**

**S6 Figure**

**Feature classification summary across Boruta bootstraps (N = 1,000) with Random Forest importance.**

**S7 Figure**

**Feature classification summary across Boruta bootstraps (N = 1,000) with lasso importance.**

**S8 Figure**

**Feature classification summary across Boruta bootstraps (N = 1,000) with SVM importance.**

**S9 Figure**

**Feature classification summary across Boruta bootstraps (N = 1,000) with XGBoost importance.**

**S10 Figure**

**Recurrent important gene features from Boruta bootstraps with 5 importance metrics, showing genes confirmed in 1/1,000 bootstraps (A), or at least 5% or 50/1,000 bootstraps (B).**

**S1 Table**

**Summary descriptive statistics table by groups of binary risk group (BRG, Low = 0, Not-low = 1).**

**S2 Table**

**Differentially expressed genes (DEGs) from training set comparison (binary risk group, sBRG 0 vs 1).**

**S3 Table**

**Standardized output table showing gene-wise importance across 4 algorithms. Ensembl gene ID (column 1), gene synbol (2), lasso beta coefficients (3:5), random forest importance (6:8), XGBoost importance (9:11), and SVM weights (12:15).**

**S4 Table**

**Manifest for data download.**

## Acknowledgments

We thank organizers and participants of NCBI Hackathons for supporting this project with feedback and input from its inception.

