## Supplemental Figures for "Consensus Machine Learning for Gene Target Selection in Pediatric AML Risk"

### Slide 1
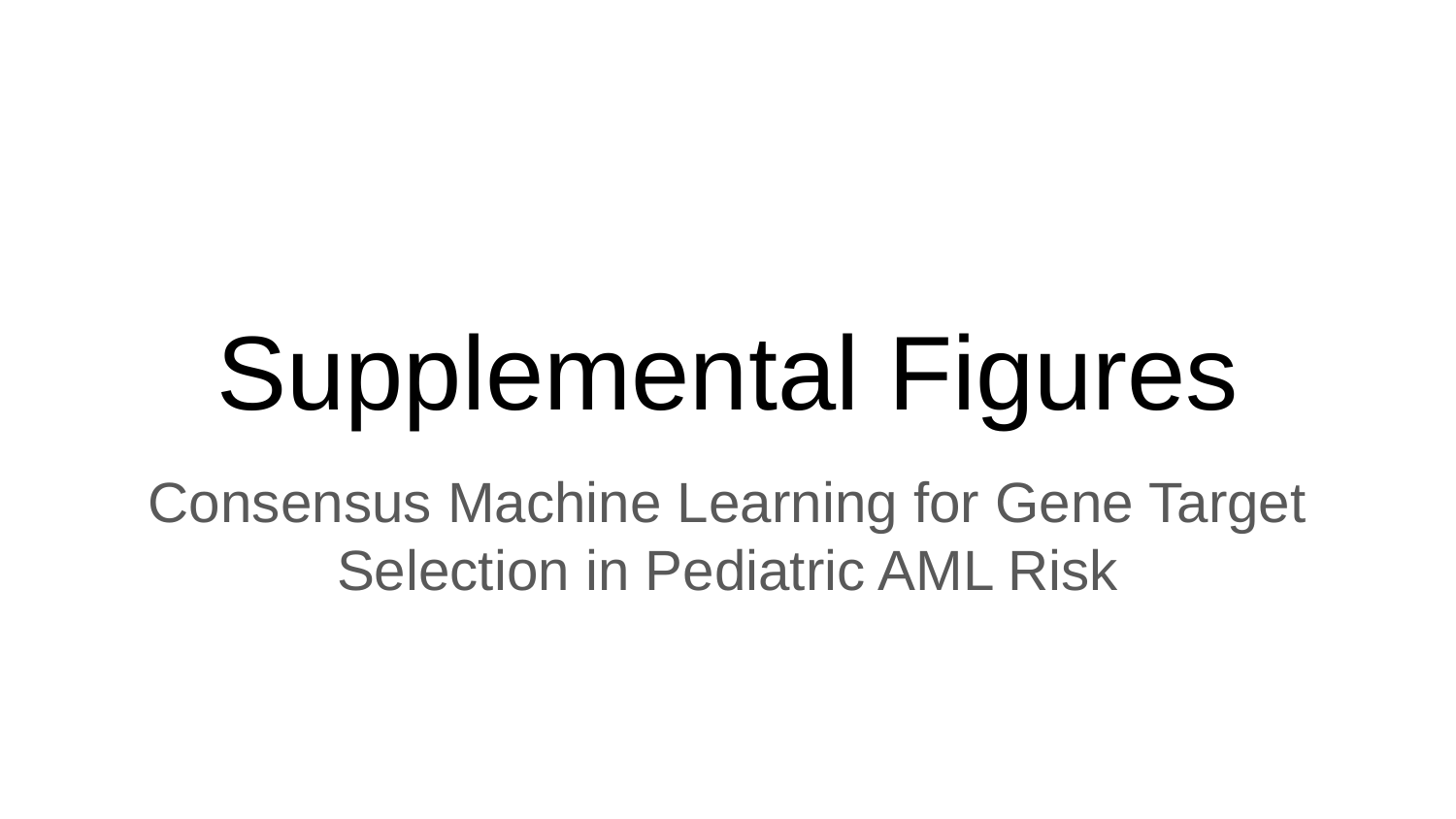

#### Supplemental Figures
Consensus Machine Learning for Gene Target Selection in Pediatric AML Risk

### Slide 2
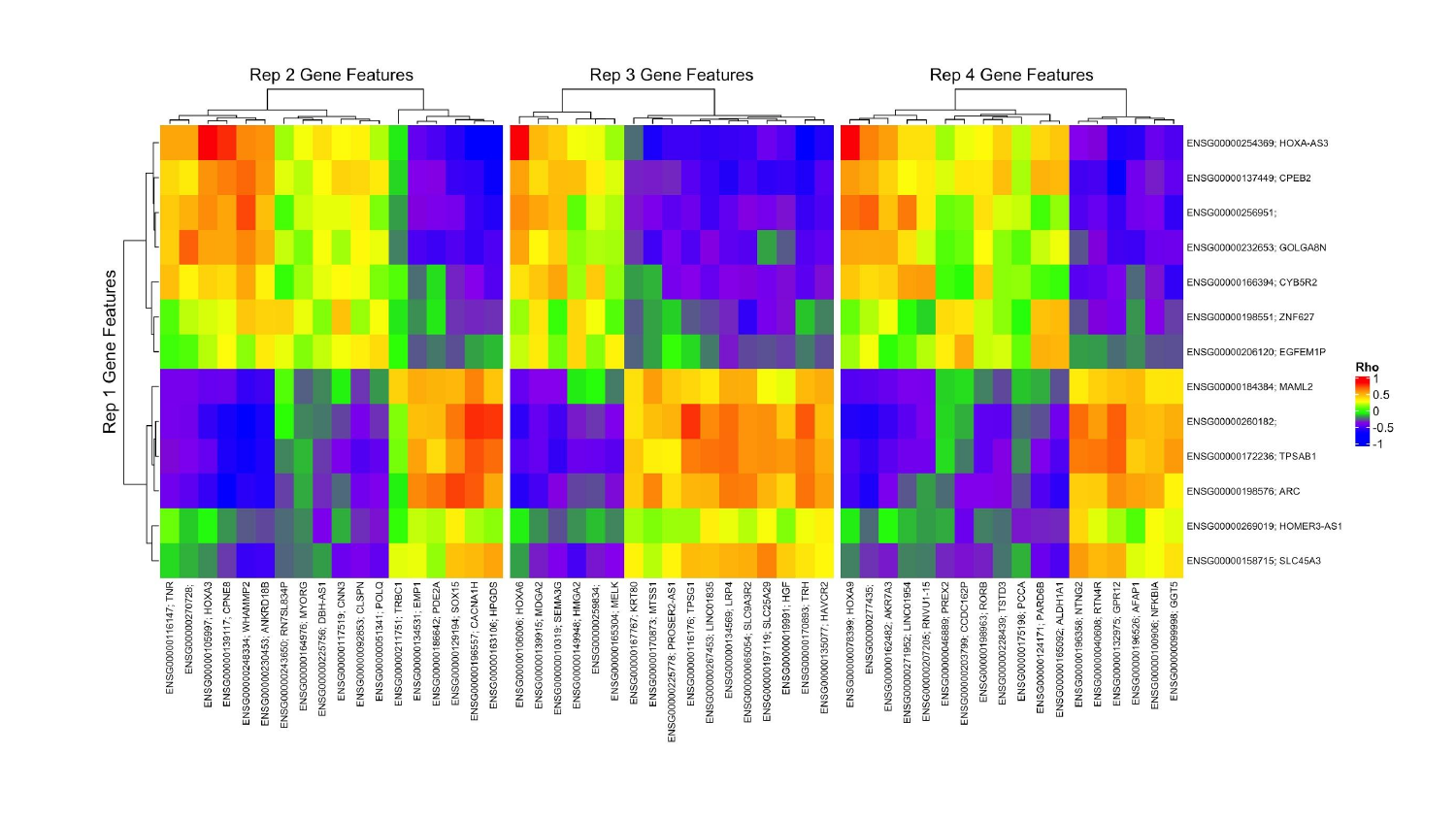

### Slide 3
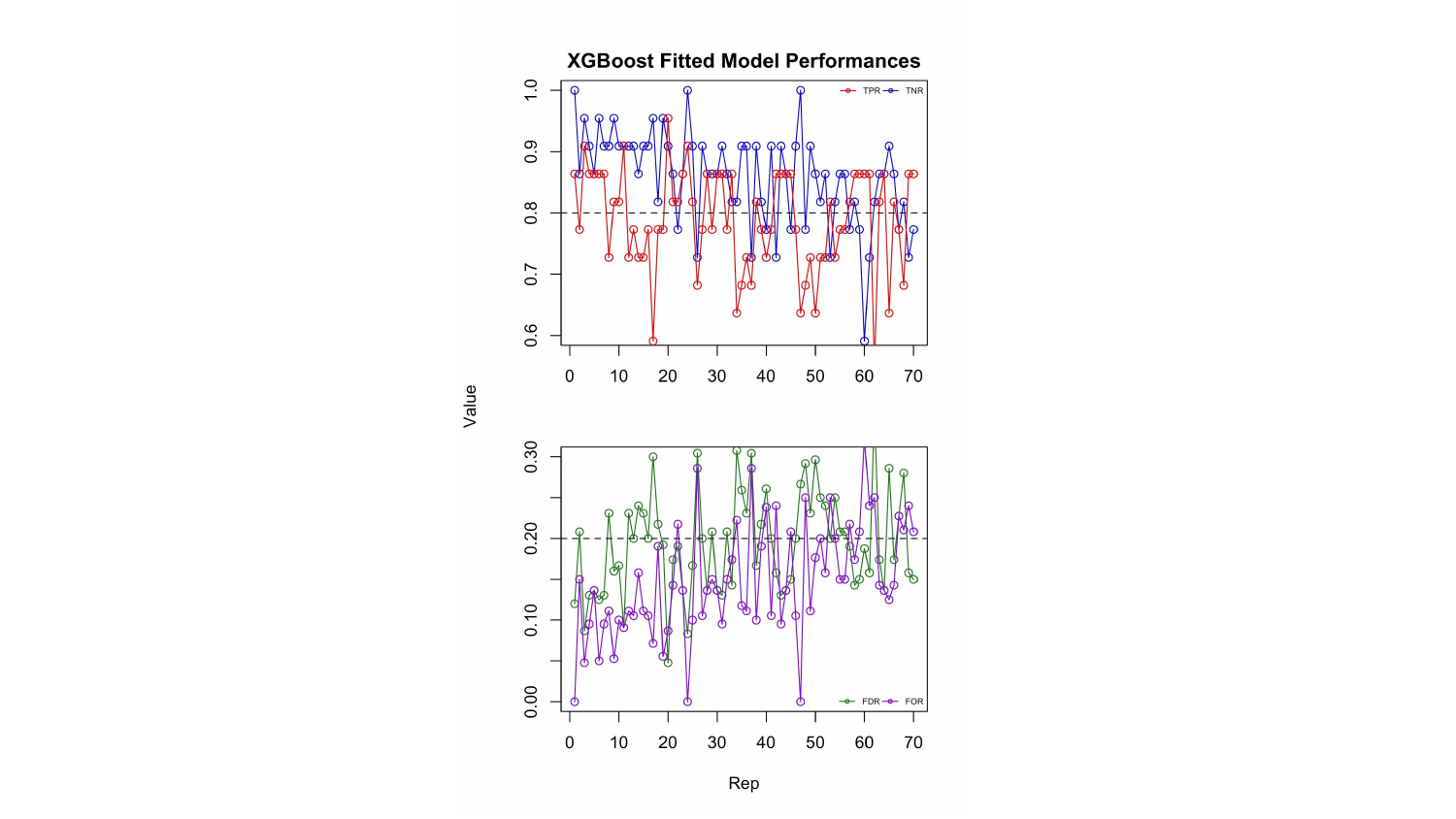

### Slide 4
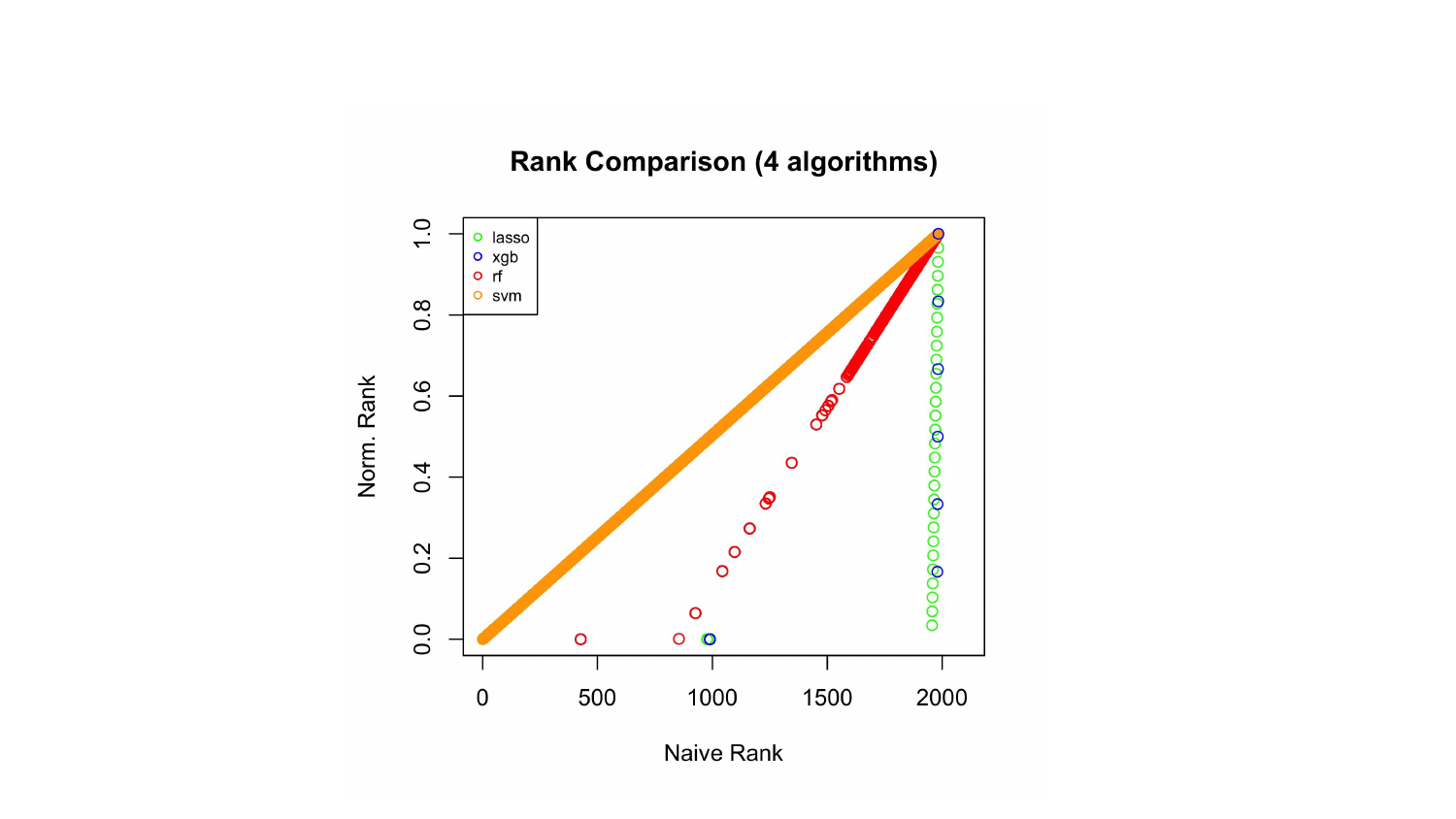

### Slide 5
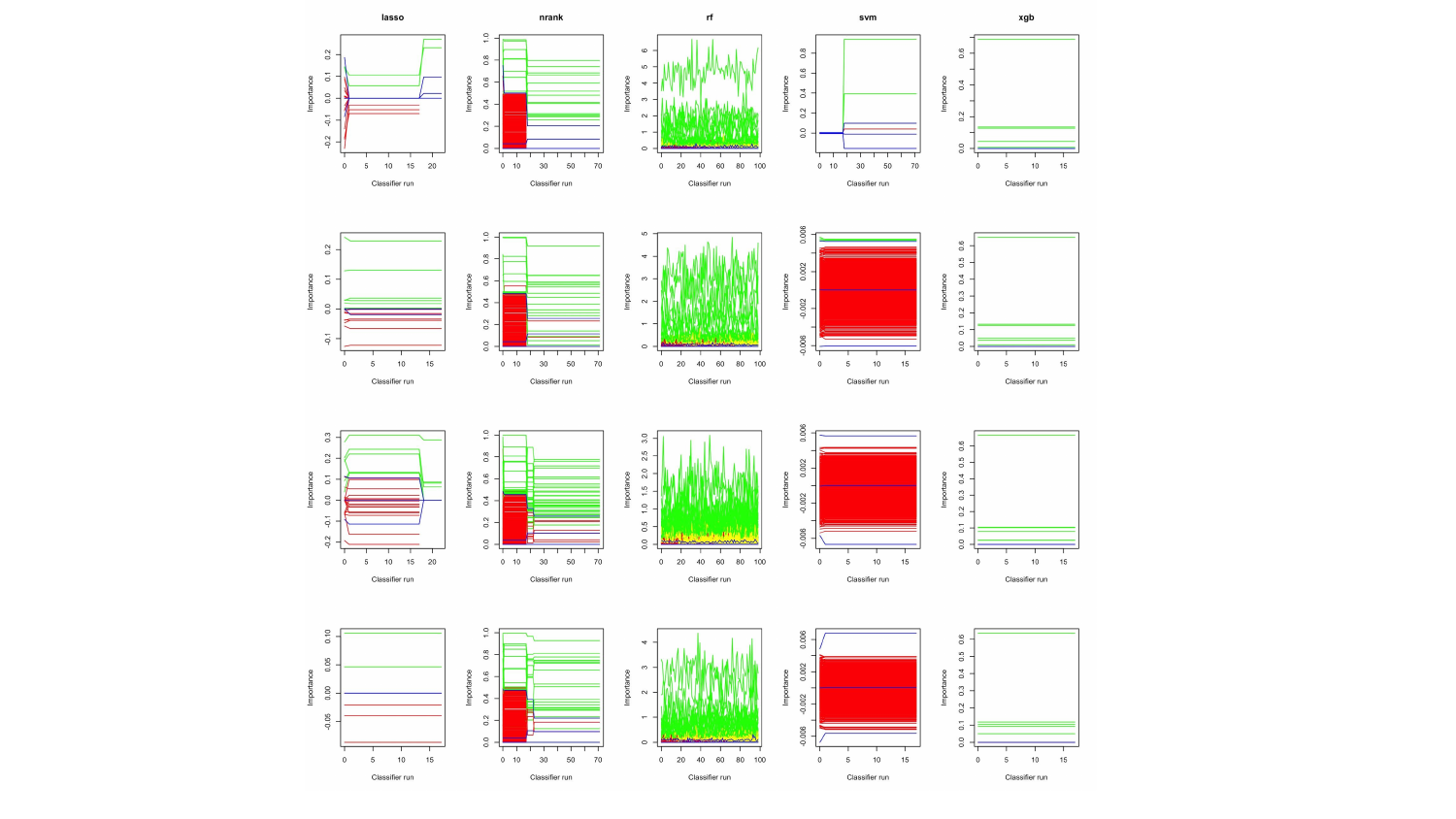

### Slide 6
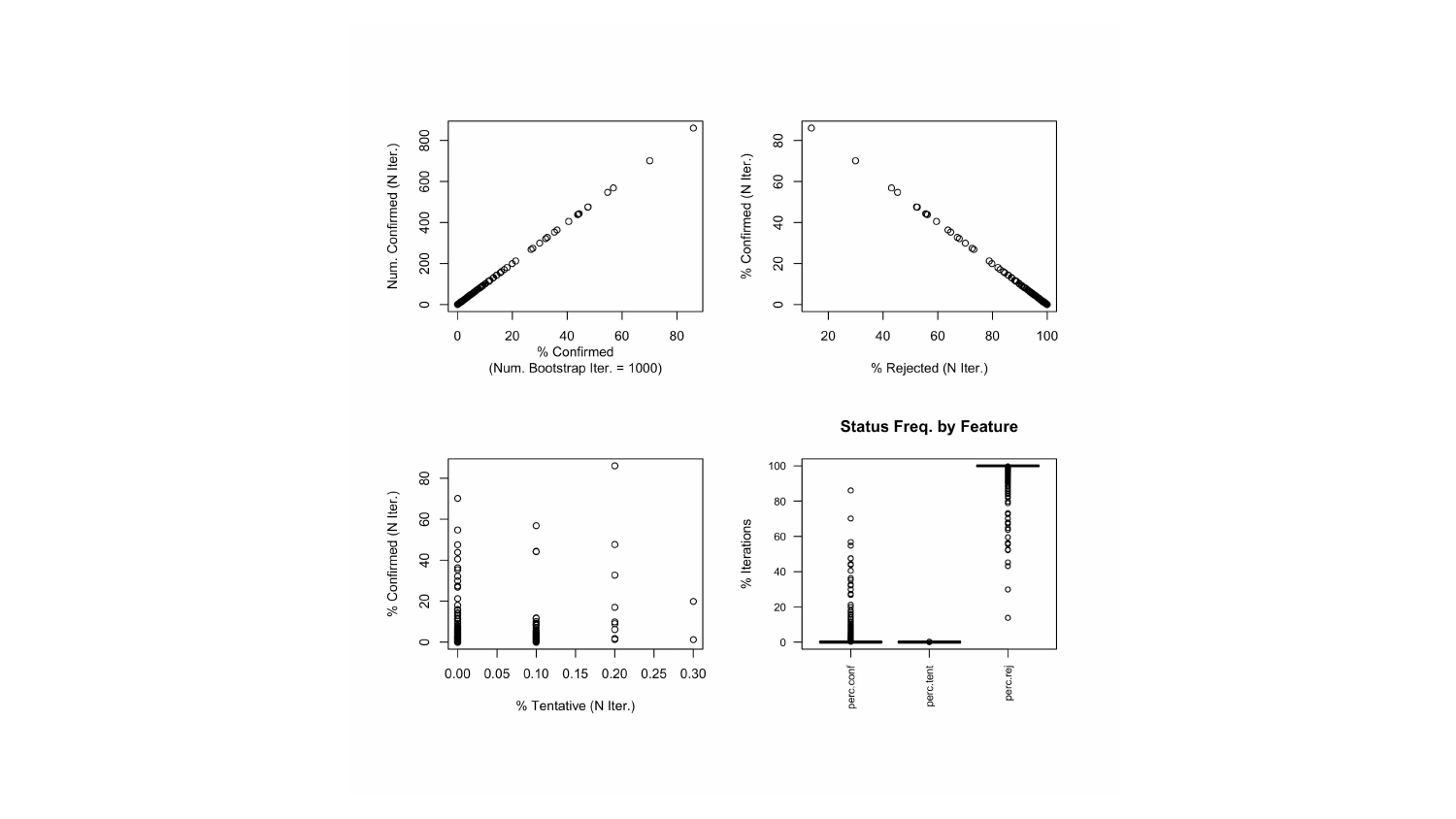

### Slide 7
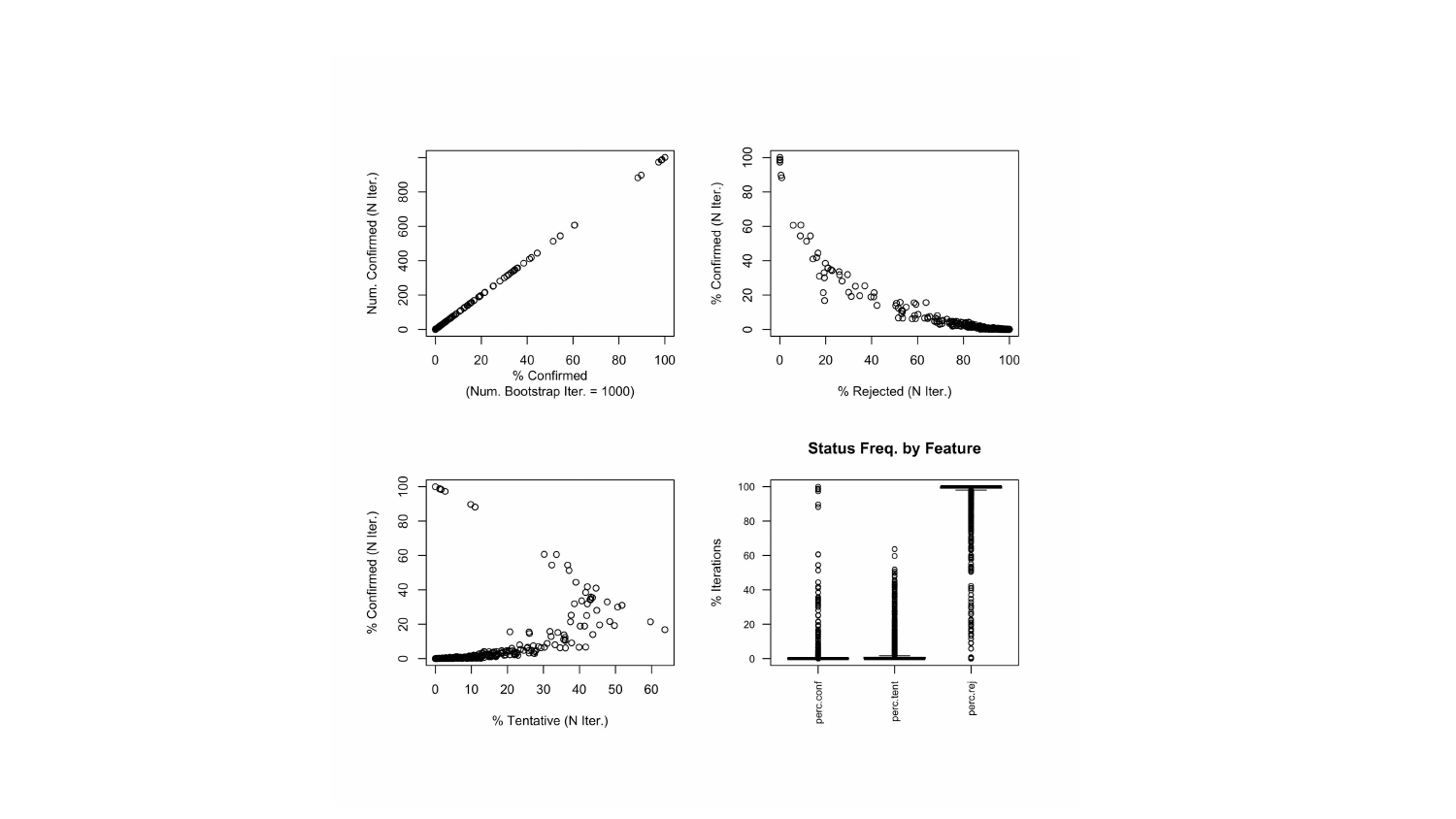

### Slide 8
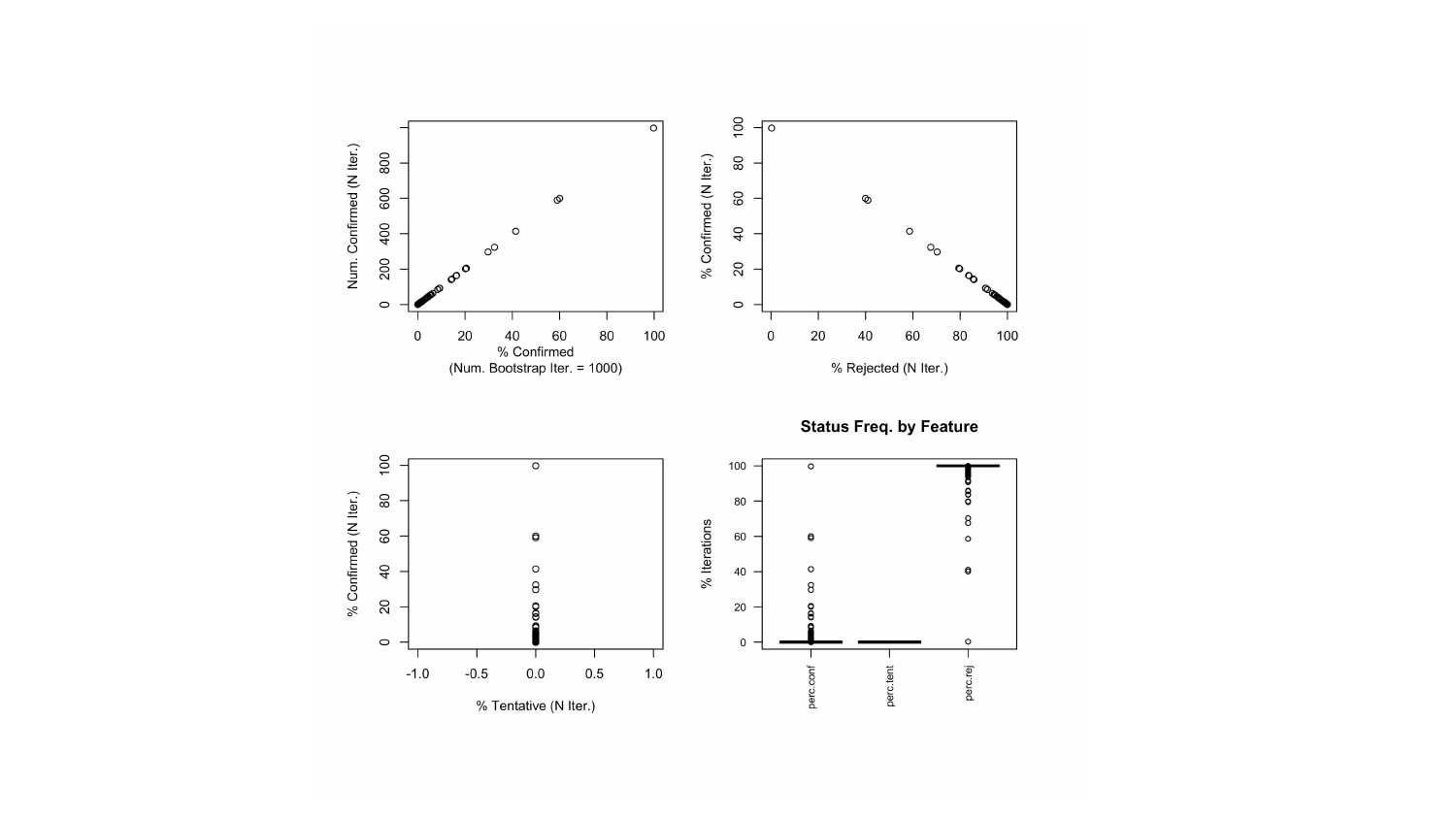

### Slide 9
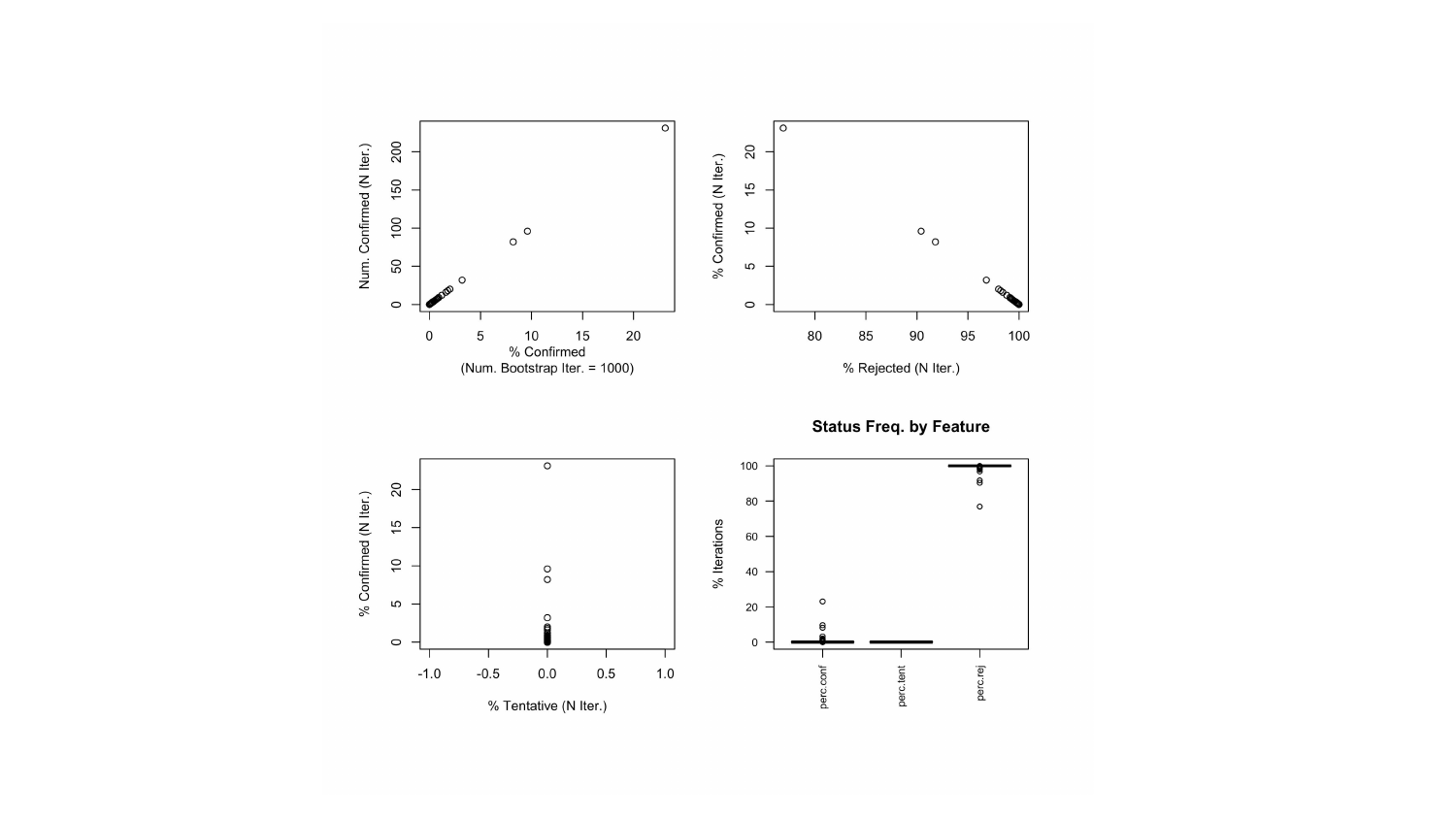

### Slide 10
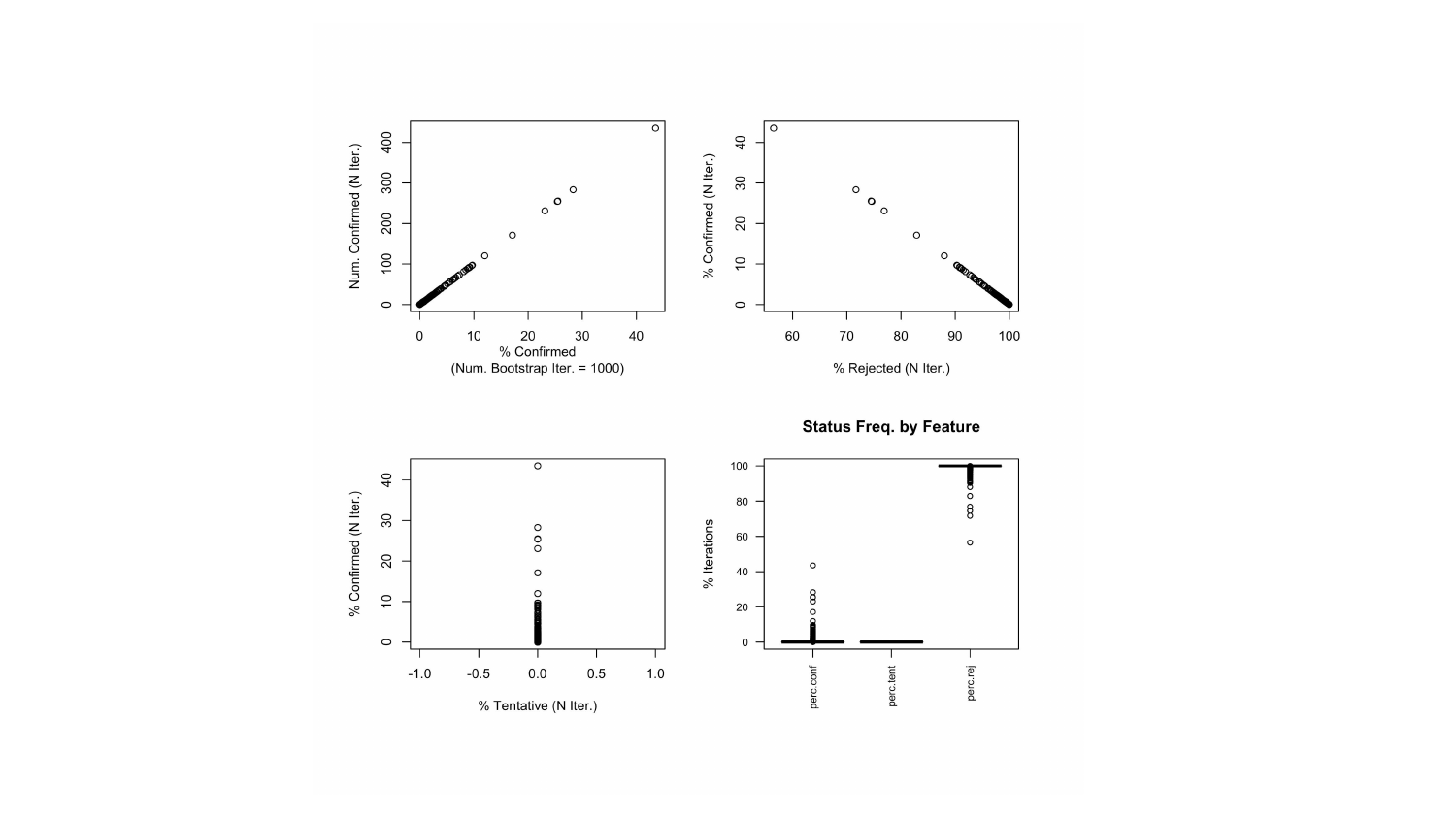

### Slide 11
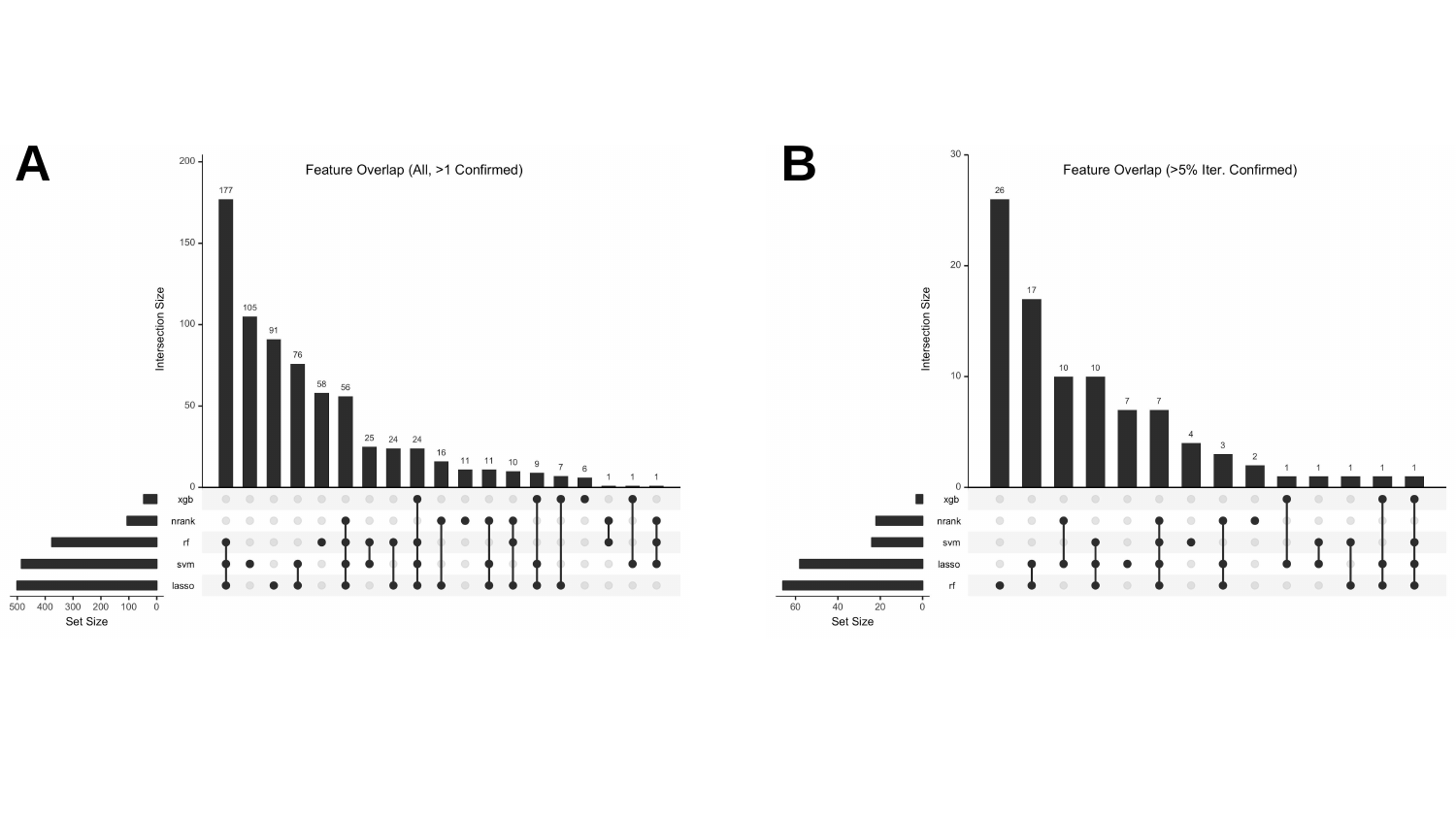

A
B
