## Supplemental Figures PDF for "Consensus Machine Learning for Gene Target Selection in Pediatric AML Risk"

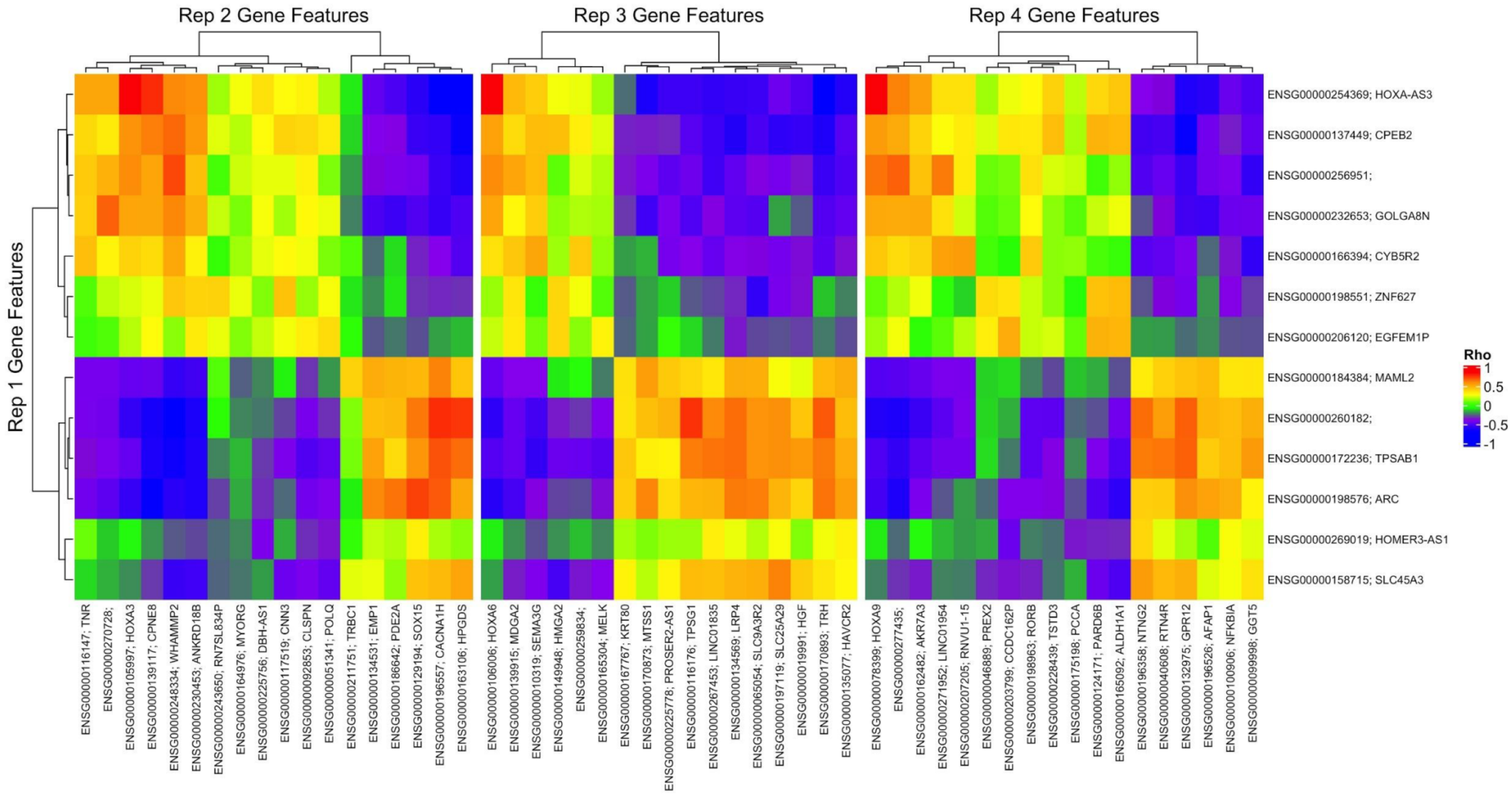

Rep 2 Gene Features
Rep 3 Gene Features
Rep 4 Gene Features
Rep 1 Gene Features
Rho
1
0.5
0
-0.5
-1
ENSG00000254369; HOXA-AS3
ENSG00000137449; CPEB2
ENSG00000256951;
ENSG00000232653; GOLGA8N
ENSG00000166394; CYB5R2
ENSG00000198551; ZNF627
ENSG00000206120; EGFEM1P
ENSG00000184384; MAML2
ENSG00000260182;
ENSG00000172236; TPSAB1
ENSG00000198576; ARC
ENSG00000269019; HOMER3-AS1
ENSG00000158715; SLC45A3
ENSG00000116147; TNR
ENSG00000270728;
ENSG00000105997; HOXA3
ENSG00000139117; CPNE8
ENSG00000248334; WHAMMP2
ENSG00000230453; ANKRD18B
ENSG00000243650; RN7SL834P
ENSG00000164976; MYORG
ENSG00000225756; DBH-AS1
ENSG00000117519; CNN3
ENSG00000092853; CLSPN
ENSG00000051341; POLQ
ENSG00000211751; TRBC1
ENSG00000134531; EMP1
ENSG00000186642; PDE2A
ENSG00000129194; SOX15
ENSG00000196557; CACNA1H
ENSG00000163106; HPGDS
ENSG00000106006; HOXA6
ENSG00000139915; MDGA2
ENSG00000010319; SEMA3G
ENSG00000149948; HMGA2
ENSG00000259834;
ENSG00000165304; MELK
ENSG00000167767; KRT80
ENSG00000170873; MTSS1
ENSG00000225778; PROSER2-AS1
ENSG00000116176; TPSG1
ENSG00000267453; LINC01835
ENSG00000134569; LRP4
ENSG00000065054; SLC9A3R2
ENSG00000197119; SLC25A29
ENSG00000019991; HGF
ENSG00000170893; TRH
ENSG00000135077; HAVCR2
ENSG00000078399; HOXA9
ENSG00000277435;
ENSG00000162482; AKR7A3
ENSG00000271952; LINC01954
ENSG00000207205; RNVU1-15
ENSG00000046889; PREX2
ENSG00000203799; CCDC162P
ENSG00000198963; RORB
ENSG00000228439; TSTD3
ENSG00000175198; PCCA
ENSG00000124171; PARD6B
ENSG00000165092; ALDH1A1
ENSG00000196358; NTNG2
ENSG00000040608; RTN4R
ENSG00000132975; GPR12
ENSG00000196526; AFAP1
ENSG00000100906; NFKBIA
ENSG00000099998; GGT5

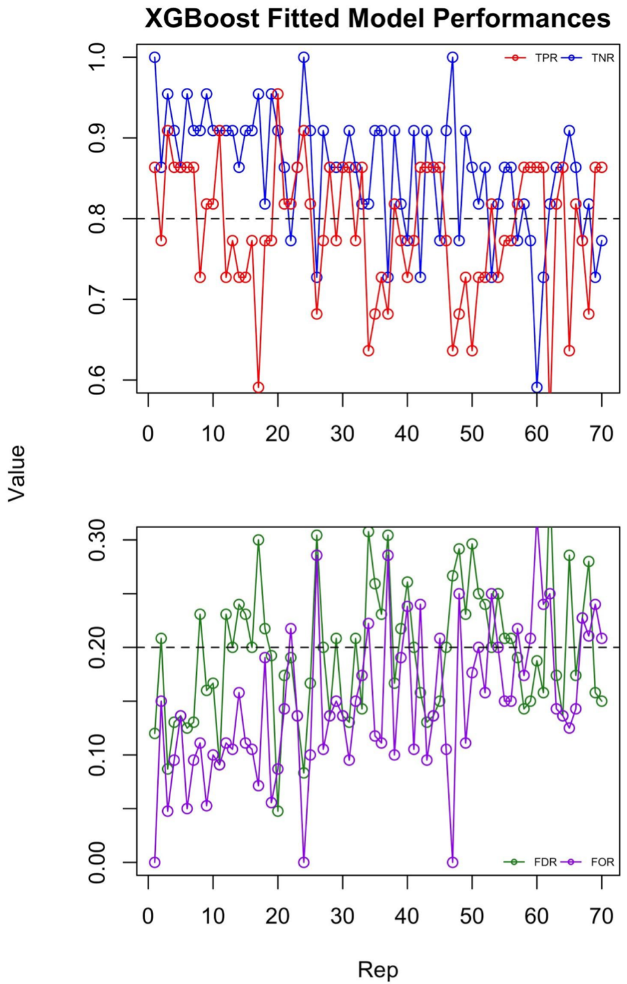
XGBoost Fitted Model Performances
TPR
TNR
FDR
FOR
Value
Rep
0.6
0.7
0.8
0.9
1.0
0.00
0.10
0.20
0.30
0
10
20
30
40
50
60
70

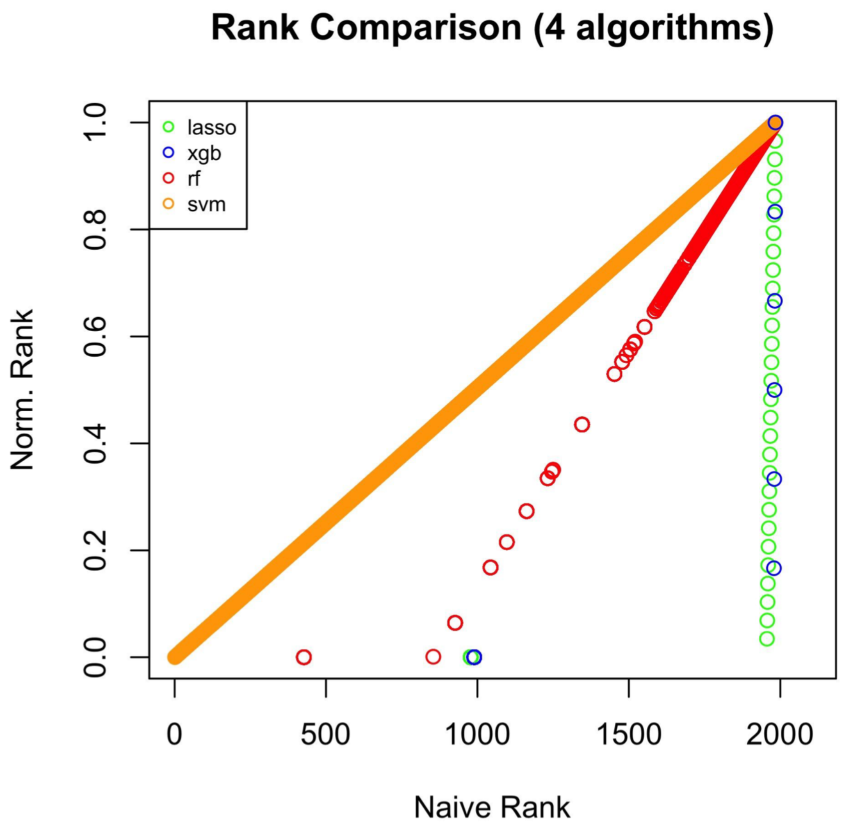
Rank Comparison (4 algorithms)
lasso
xgb
rf
svm
Norm. Rank
Naive Rank
0.0
0.2
0.4
0.6
0.8
1.0
0
500
1000
1500
2000

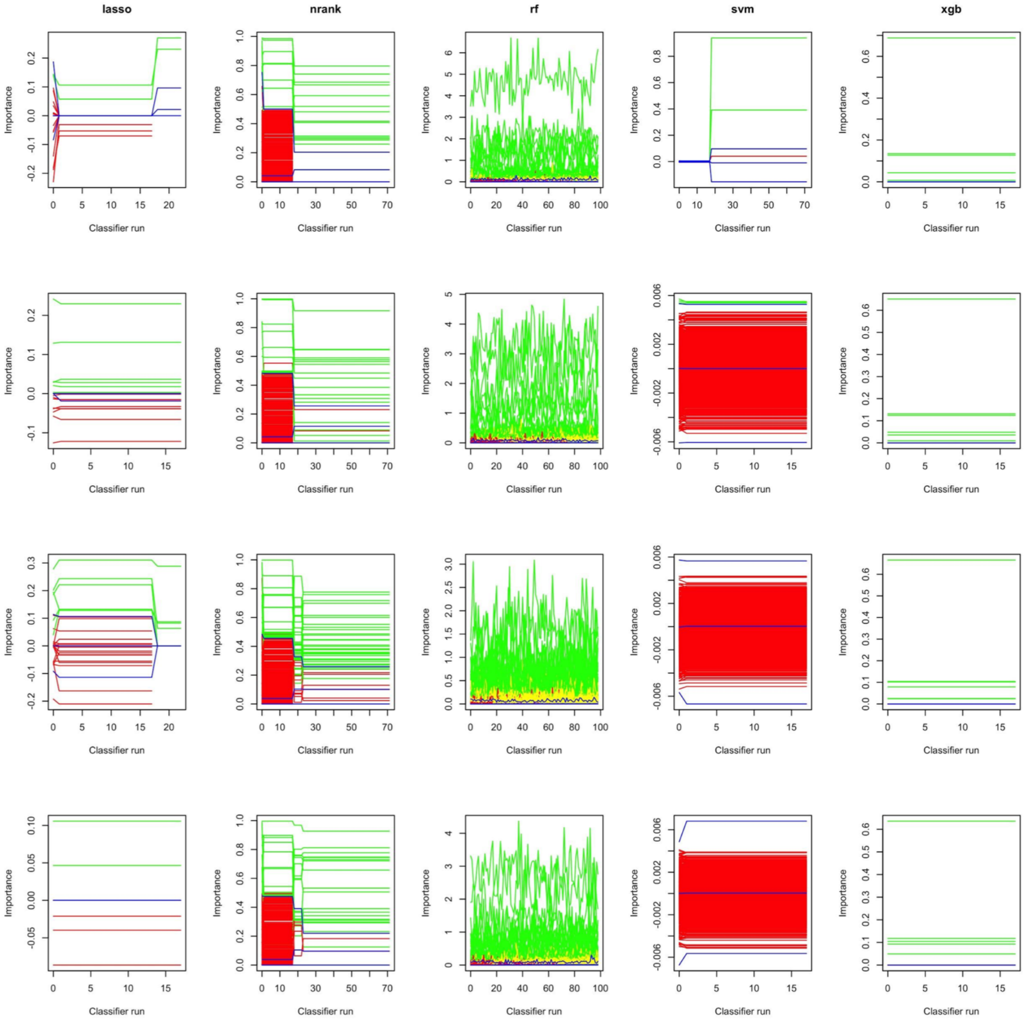
lasso
nrank
rf
svm
xgb
Importance
Classifier run

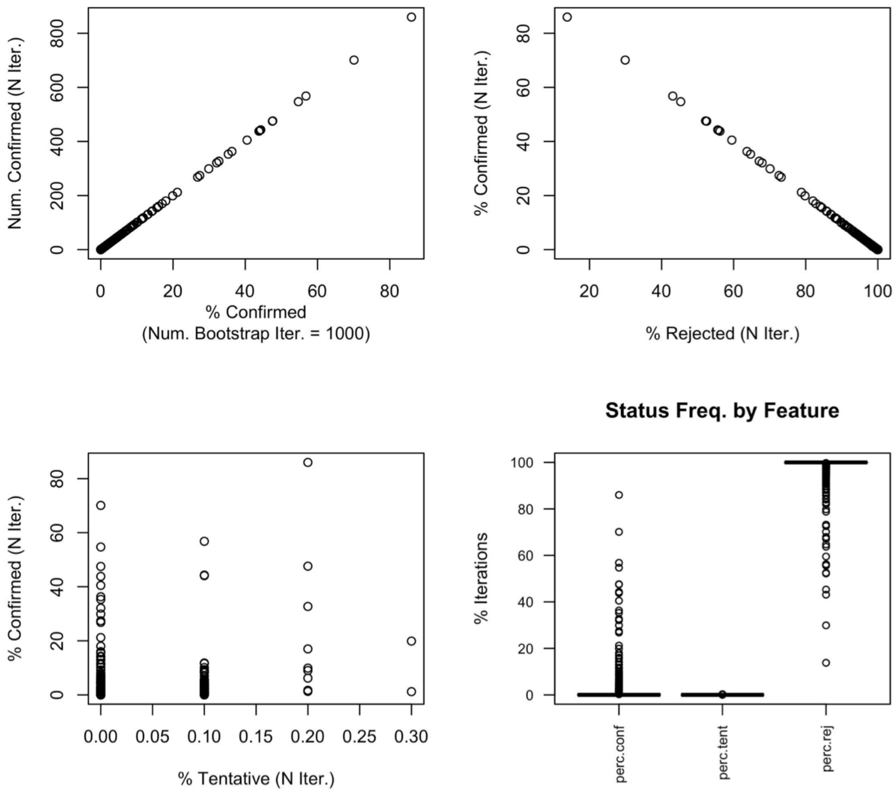
Num. Confirmed (N Iter.)
% Confirmed
(Num. Bootstrap Iter. = 1000)
% Confirmed (N Iter.)
% Rejected (N Iter.)
% Confirmed (N Iter.)
% Tentative (N Iter.)
Status Freq. by Feature
% Iterations
perc.conf
perc.tent
perc.rej

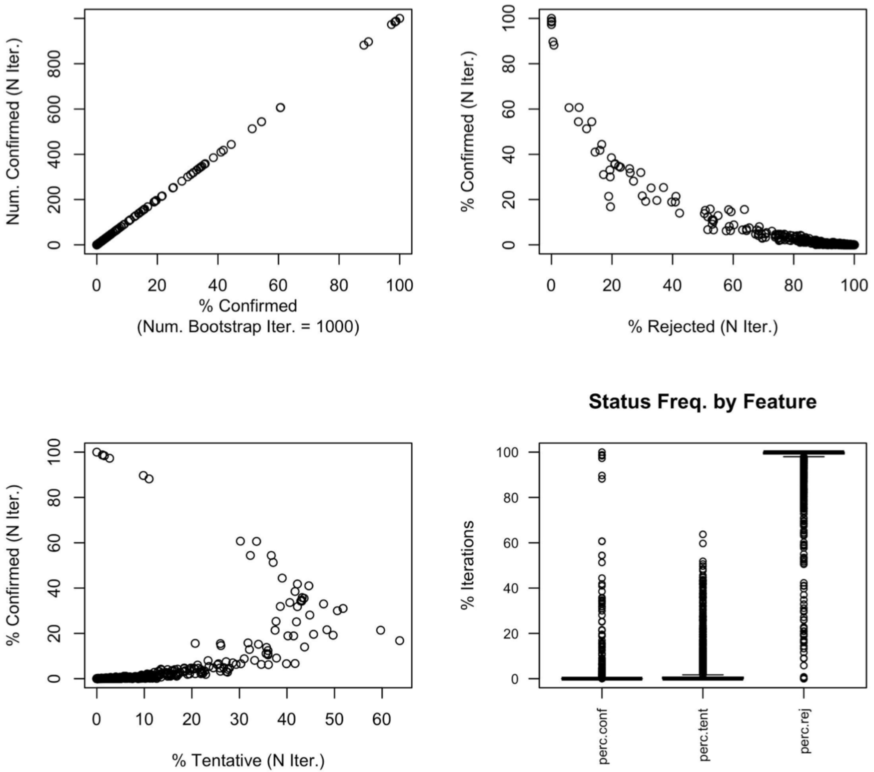
Num. Confirmed (N Iter.)
% Confirmed
(Num. Bootstrap Iter. = 1000)
% Confirmed (N Iter.)
% Rejected (N Iter.)
% Confirmed (N Iter.)
% Tentative (N Iter.)
Status Freq. by Feature
% Iterations
perc.conf
perc.tent
perc.rej

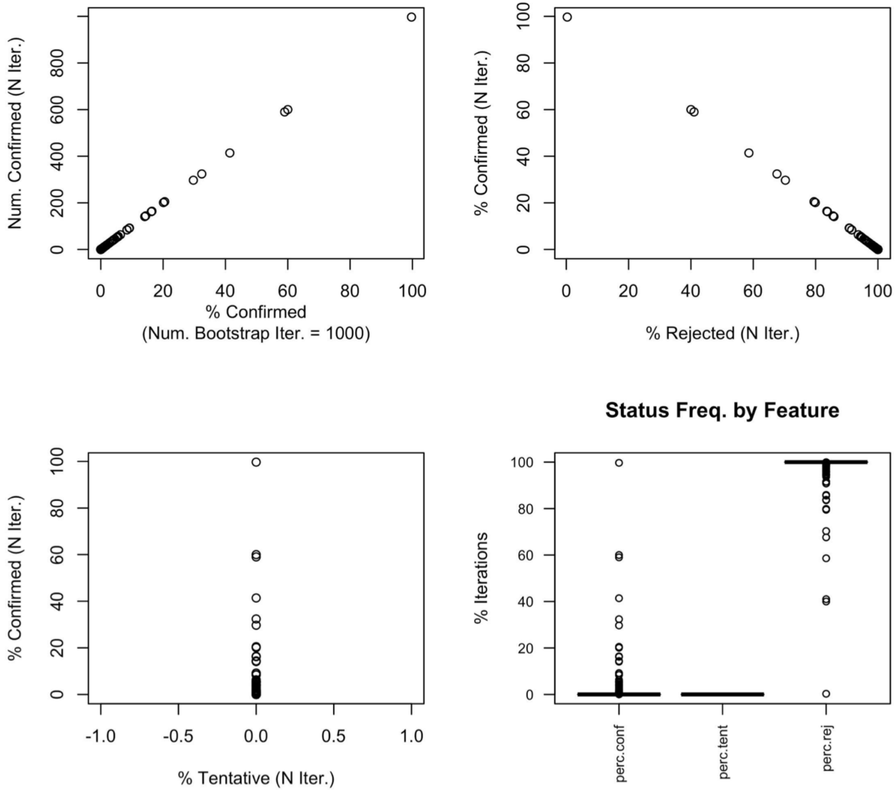
Num. Confirmed (N Iter.)
% Confirmed
(Num. Bootstrap Iter. = 1000)
% Confirmed (N Iter.)
% Rejected (N Iter.)
Status Freq. by Feature
% Confirmed (N Iter.)
% Tentative (N Iter.)
% Iterations
perc.conf
perc.tent
perc.rej

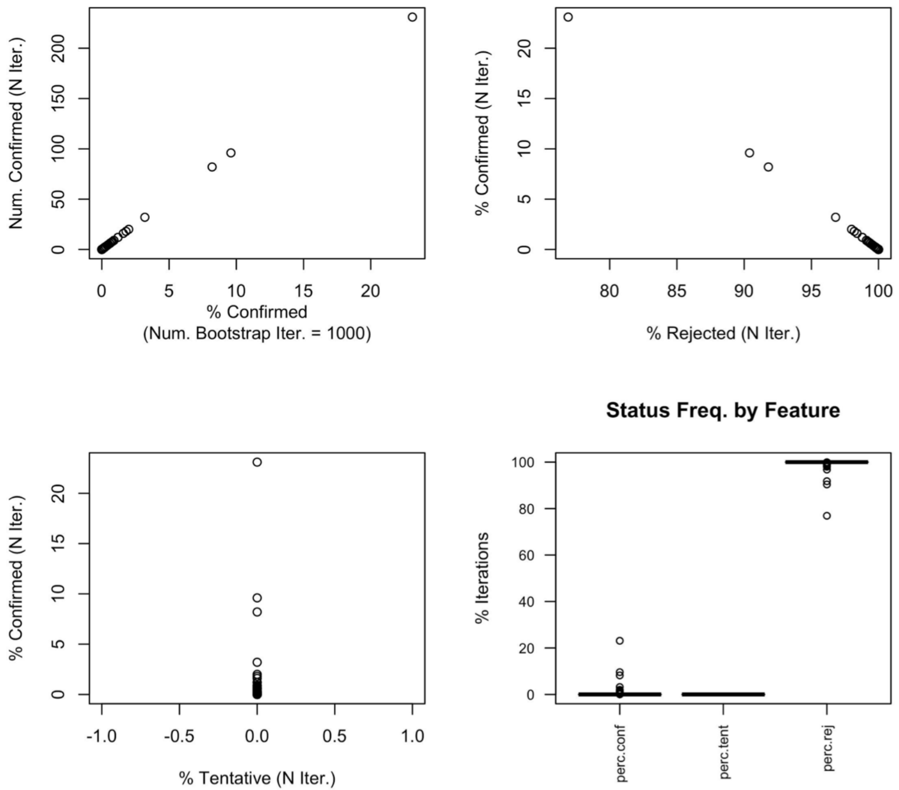
Num. Confirmed (N Iter.)
% Confirmed
(Num. Bootstrap Iter. = 1000)
% Confirmed (N Iter.)
% Rejected (N Iter.)
Status Freq. by Feature
% Confirmed (N Iter.)
% Tentative (N Iter.)
% Iterations
perc.conf
perc.tent
perc.rej

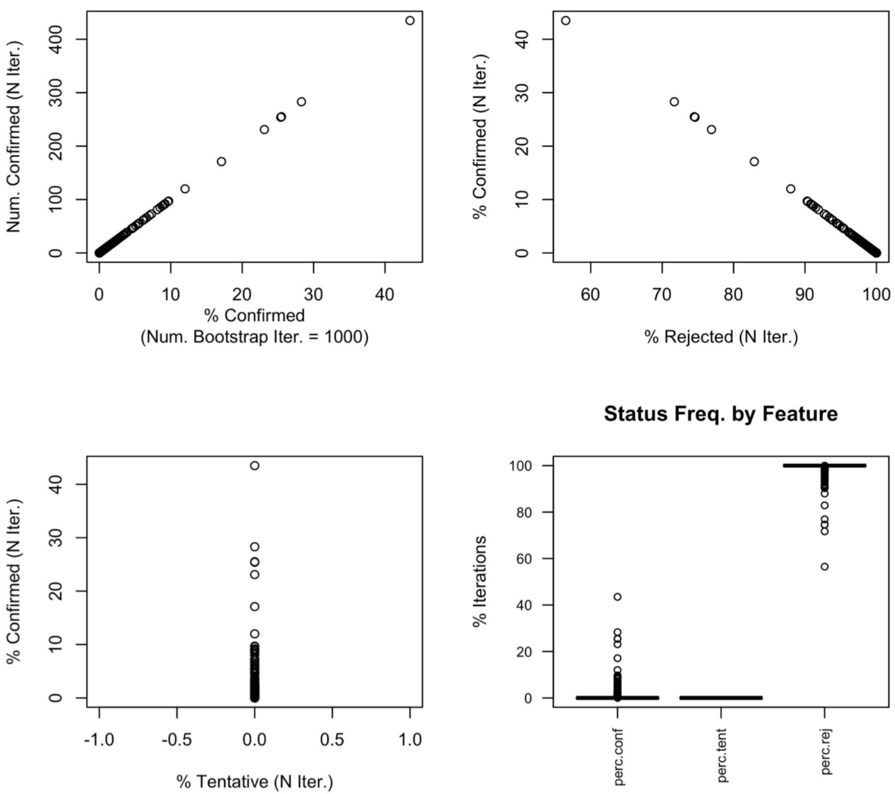
Num. Confirmed (N Iter.)
% Confirmed
(Num. Bootstrap Iter. = 1000)
% Confirmed (N Iter.)
% Rejected (N Iter.)
Status Freq. by Feature
% Confirmed (N Iter.)
% Tentative (N Iter.)
% Iterations
perc.conf
perc.tent
perc.rej

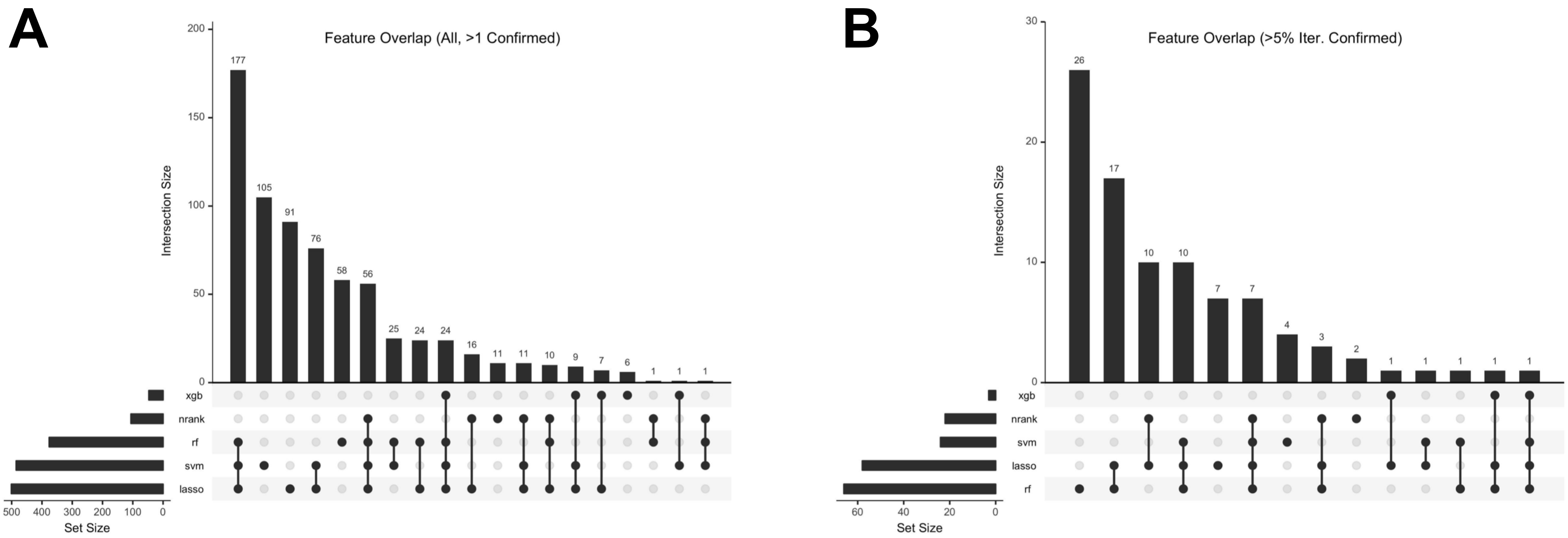
A
Feature Overlap (All, >1 Confirmed)
Intersection Size
177
105
91
76
58
56
25
24
24
16
11
11
10
9
7
6
1
1
1
200
150
100
50
0
xgb
nrank
rf
svm
lasso
500
400
300
200
100
0
Set Size
B
Feature Overlap (>5% Iter. Confirmed)
Intersection Size
26
17
10
10
7
7
4
3
2
1
1
1
1
1
30
20
10
0
xgb
nrank
svm
lasso
rf
60
40
20
0
Set Size
