## Supplementary material for "Consensus Machine Learning for Gene Target Selection in Pediatric AML Risk": Boruta Methods Notebook

### Bootstrapping Boruta Permutations

This notebook describes implementation of the Boruta permutations method to identify consensus features from the TARGET pediatric AMl dataset.

#### 1. Methods

We applied Boruta as implemented in the Boruta R package (v.6.0.0). We implemented the Boruta permuataions method with a consensus rank importance metric, and conducted bootstraps of data subset selection (2/3rds of data, to simulate iteratively redraw of training sample subset). Each Boruta iteration was allowed to run for up to 100 steps, over which the algorithm designated "confirmed", "tentative", and "rejected" features.

Hyperparameter sets used for algorithm with Boruta permuatations were informed by preliminary investigation of model performances across hyperparameter sets for each algorithm. The initial defined training and test data subsets, as used to measure differential gene expression, were used in hyperparameter optimization and to derive the rank-based consensus importance measure as described below.

##### 1A. Consensus Importance ('nrank')

To calculate a consensus importance metric, we define a method to take the median normalized absolute rank of features from across 4 algorithms (lasso, XGBoost, Random Forest, and SVM).

Steps to calculate this normalized median importance include:

1. Calculate algorithm-specific importances.
2. Take absolute importances from (1).
3. Calculate naive rank (index-based) of importances from (2).
4. Calcualte normalized rank from (3).
5. Calculate median normalized rank and return.

We settled on this strategy due to properties of Boruta and feature selection. For comparison, a standard of Boruta using Random Forest importance shows oscillation of shadow feature importances, gradual exclusion of rejected features, and retention of important features (Figure 1)

By contrast, using a naive median rank as our consensus importance yields a static pattern of shadow feature importance across permutations (Figure 2).

The static nature of naive rank is almost certainly due to immediate fixation of feature consensus importance from the first iteration. This is likely because naive rank is not normalized, and for penalized algorithms (XGBoost and lasso), it returns very low rank for most features. These characteristics make this metric less desriable for use in Boruta permutations, as importance estimation does not benefit from shadow feature importance and background estimation.

Using the steps described above, we calculated a normalized median rank for the consensus importance we ultimately used to identify consensus important features. A single Boruta iteration using this metric shows progressive retention of "confirmed" and "tentative" features, and loss of "rejected" features over permutations, indicating this metric does benefit from shadow feature iterations (Figure 3).

Comparing consensus importance derived from naive median rank and normalized median rank directly, there is greater variation in importance measured using the latter (Figure 4).

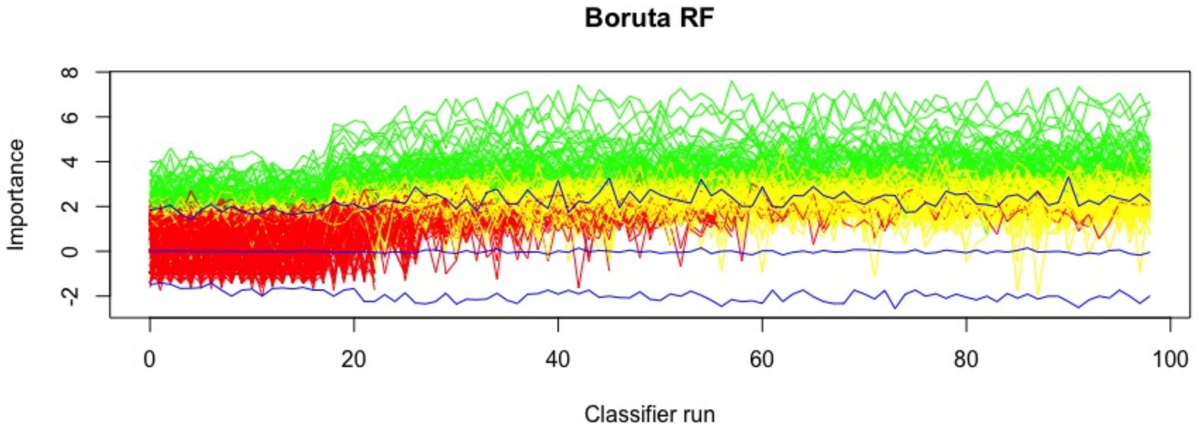


Figure 1: History of shadow feature importances over permuatations in a single Boruta iteration using Random Forest importance (green: confirmed features, red: rejected features, yellow: tentative features).

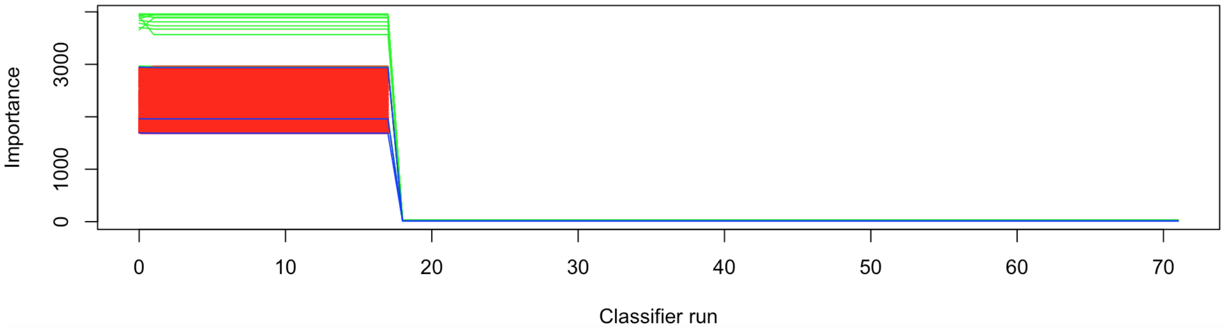


Figure 2: History of shadow feature importance over Boruta permutations using naive median rank as consensus importance (green: confirmed; red: rejected).

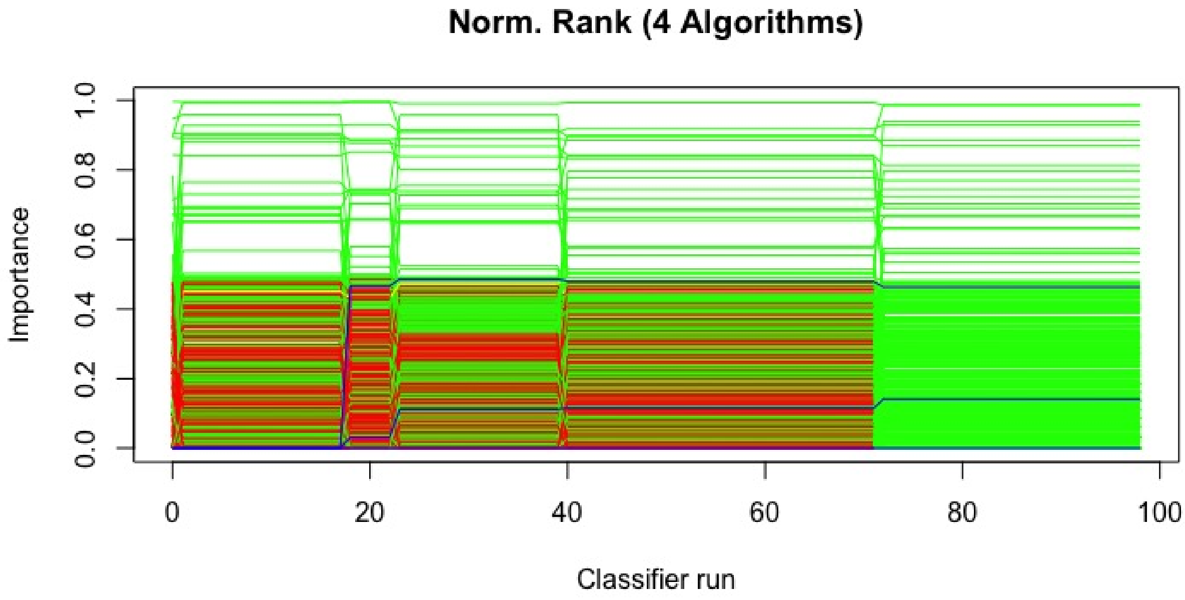


Figure 3: History of shadow feature importance with normalized median rank used to calculate consensus importance (green: confirmed; red: rejected; yellow: tentative)

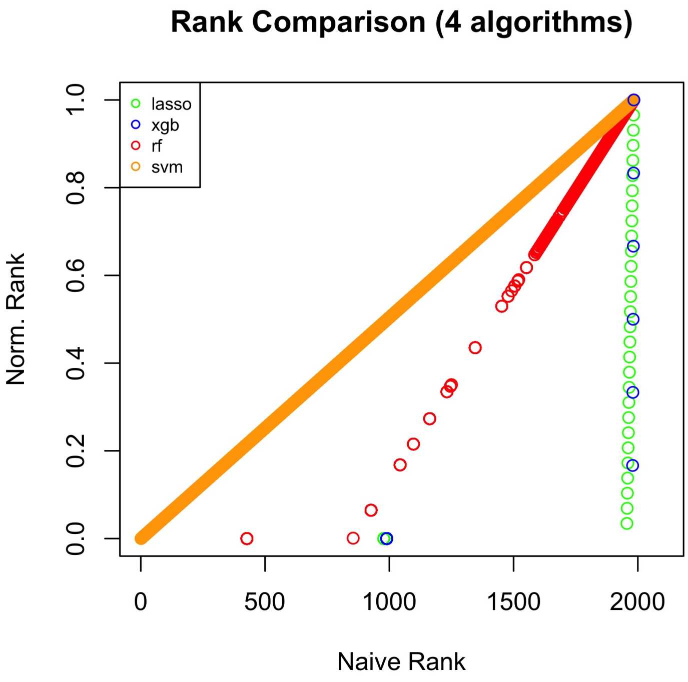


Figure 4: Comparison of median rank measures (naive vs. normalized) across 4 algorithms (lasso: green, XGBoost: blue, Random Forest: orange, and SVM: red).

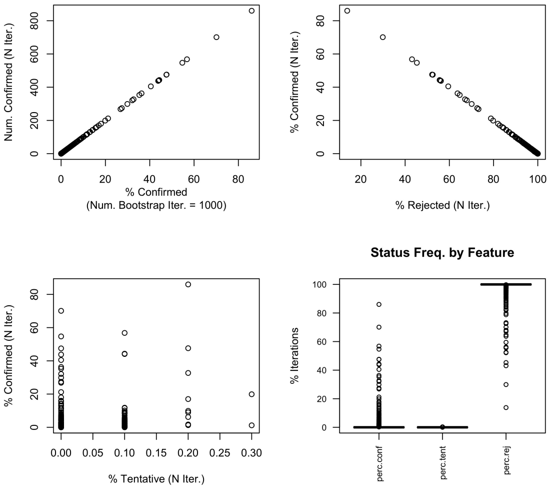


Figure 5: Feature classification summary across Boruta Bootstraps with Consensus importance (“nrank”).

#### 1B. Bootstrapping Boruta Permutations

We performed 1000 bootstraps of Boruta permutations for each of five respective importance metrics. Each bootstrap redrew two thirds of the dataset, with preservation of sample group frequency to overall frequency. This was to simulate redrawing training data subsets. The importance metrics applied included:

1. Consensus importance (normalized median absolute rank, see above).
2. Lasso importance (Beta-value coefficient).
3. XGBoost importance (feature importance).
4. Random Forest Importance (feature importance).
5. SVM Importance (feature weight).

Code for running bootstraps is described below (see Code Supplement).

### 2. Results of Boruta Permutations Bootstraps

After performing bootstraps as described, we analyzed and summarized results.

#### 2A. Summaries Across Importance Functions

We observed the following results from each bootstrap iteration employing one of the five described importance measures.

#### 2B. Feature Importance Histories Across Runs

Across bootstrap runs with varying importance metrics, we observed the following patterns of feature importnace histories from 4 bootstraps selected at random from each run.

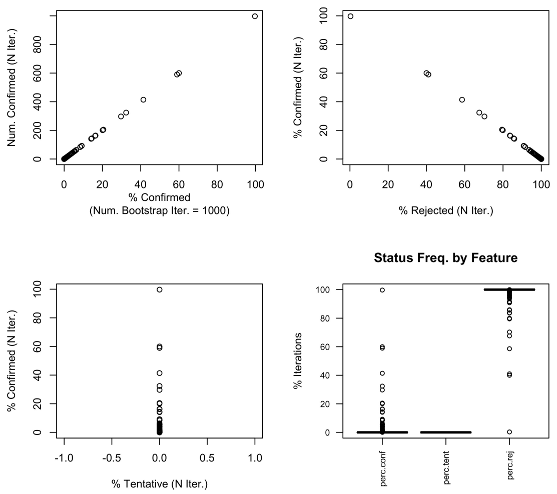


Figure 6: Feature classification summary across Boruta Bootstraps with Lasso importance.

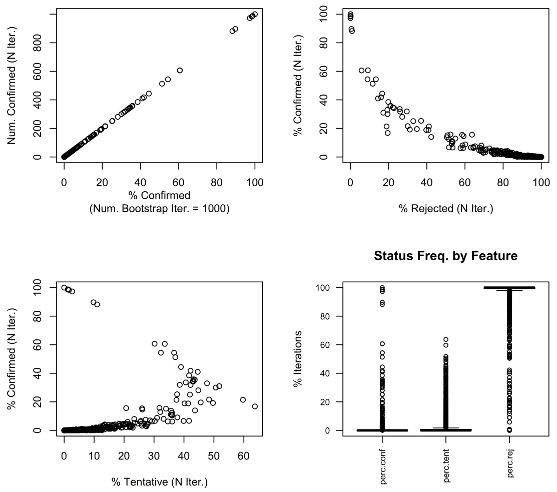


Figure 7: Feature classification summary across Boruta Bootstraps with Random Forest importance.

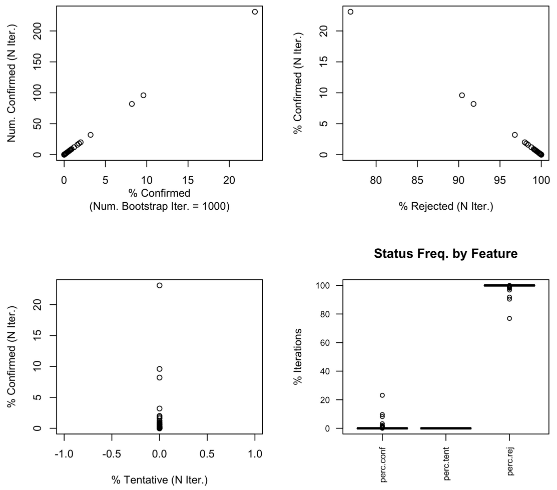


Figure 8: Feature classification summary across Boruta Bootstraps with SVM importance.

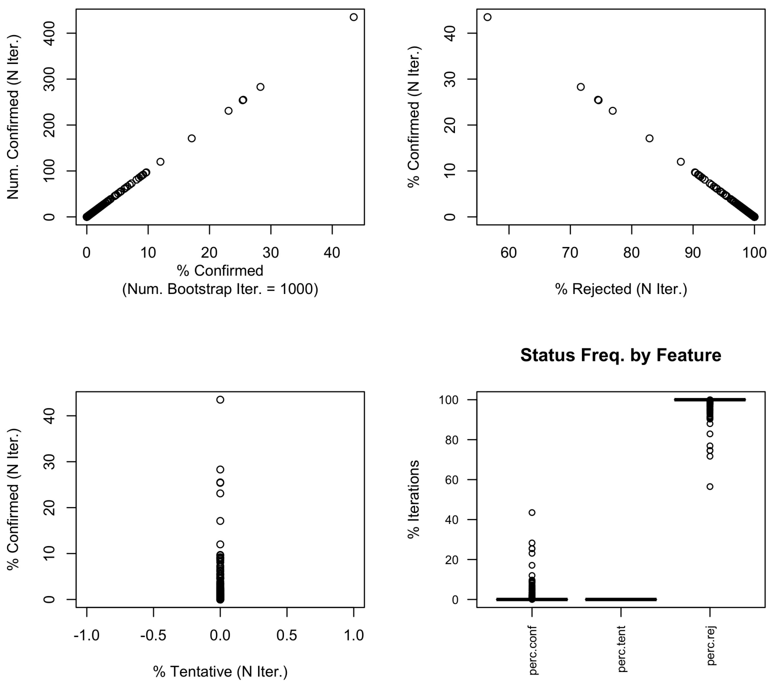


Figure 9: Feature classification summary across Boruta Bootstraps with XGBoost importance.

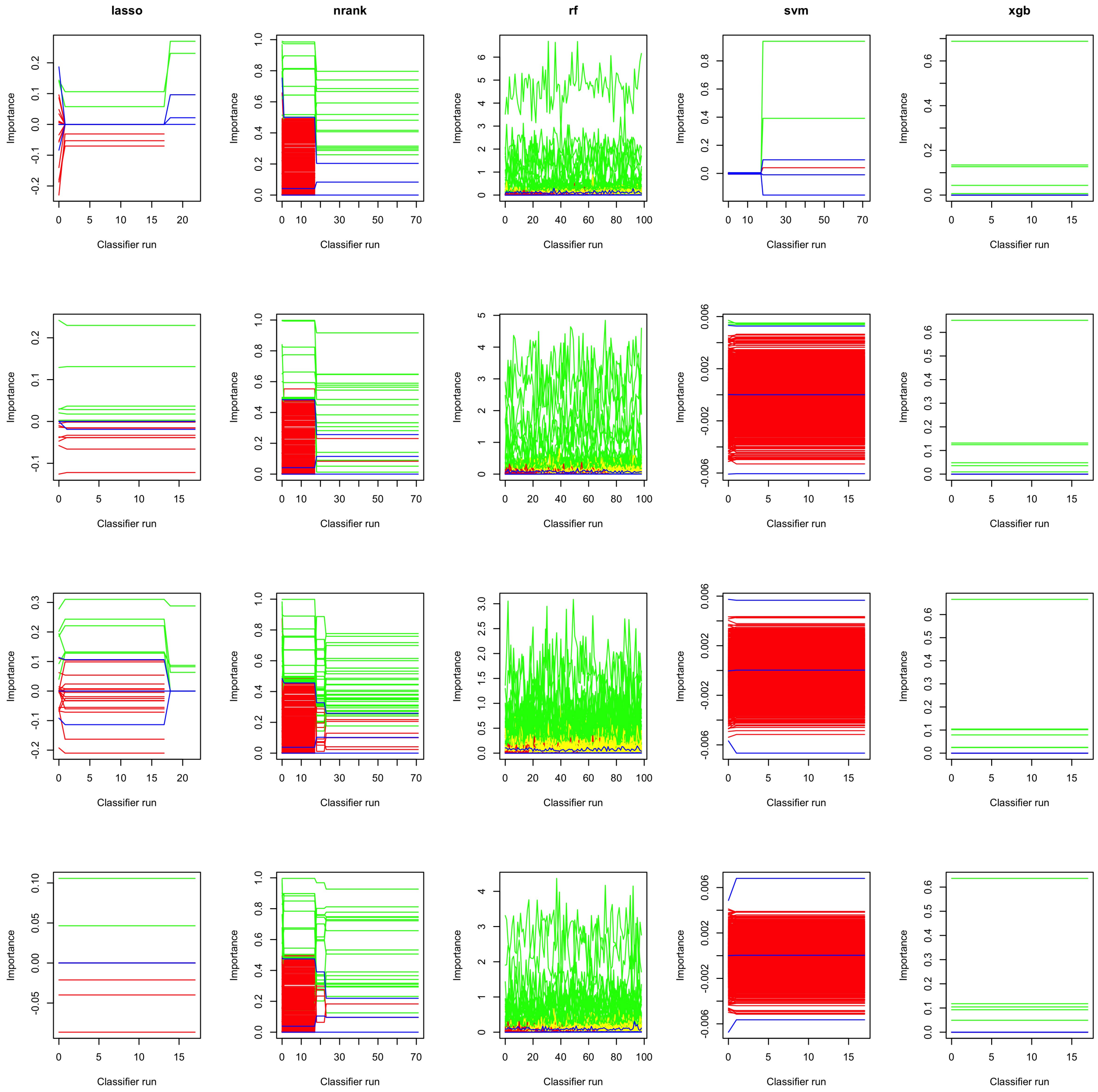


Figure 10: Feature histories across Boruta permutations, from 4 bootstraps drawn at random from each of the 5 runs (green = confirmed, red = rejected, yellow = tentative, blue lines = shadow feature max, mean, and min).

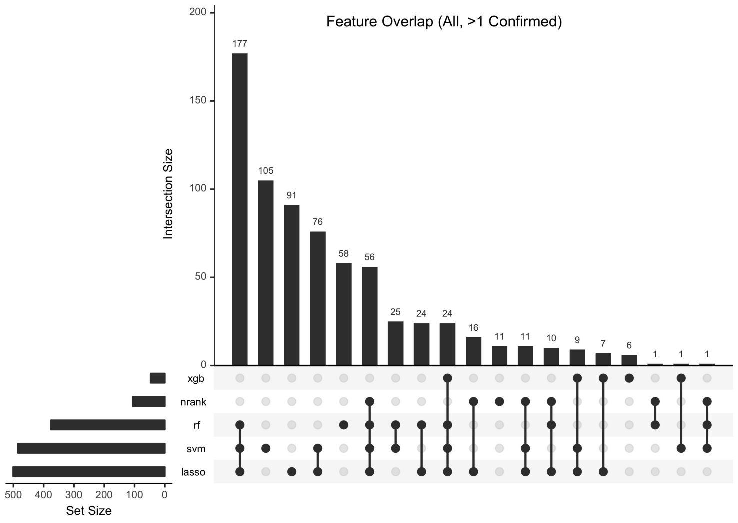


Figure 11: Feature overlap among features confirmed in at least 1 bootstrap, across bootstrap runs with 5 importance metrics.

##### 2C. Overlap of Confirmed Features

We also summarized the overlap in confirmed and recurrent confirmed features across bootstrap runs. The following upset plots show subset overlap membership for features confirmed at least once across bootstraps, greater that 5% of bootstraps, or greater than 20% of bootstraps.

#### 3. Code Supplement

This section includes supplemental code describing importance functions for Boruta permutation runs.

##### 3A. Lasso Importance

Lasso importance was defined as the beta-value coefficient assignments, or 0 for excluded genes.

```
impLasso <- function(df, classes, trainindices, seed=2019){
    # df : data frame to parse, rownames = classifier groupings, colnames = feature ids
    require(glmnet)
    require(SummarizedExperiment)
    set.seed(seed)
    var.classifier <- response <- as.character(classes)
    y <- factor(response); names(y) <- rownames(df) # response var obj
    x = df # genes of interest
    contrast <- contrasts(y)
    grid <- 10^ seq(10,-2, length=100)

    # use cross-validation on the training model.CV only for lambda
    message("performing cross-validation...")
    cv.fit <- cv.glmnet(x[trainindices,], y[trainindices], family = "binomial",
                        type.logistic="modified.Newton", standardize = FALSE,
                        lambda = grid, alpha=1, nfolds = length(trainindices), #LOOCV
```

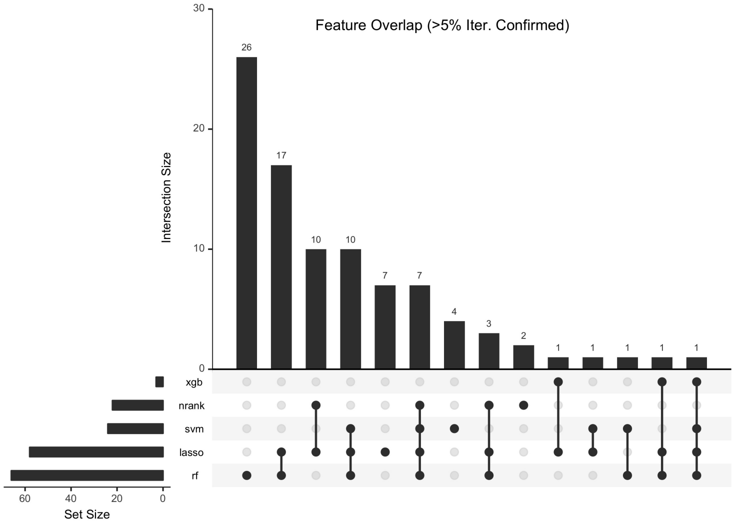


Figure 12: Feature overlap among features confirmed in at least 5% of bootstrap runs, across bootstrap runs with 5 importance metrics

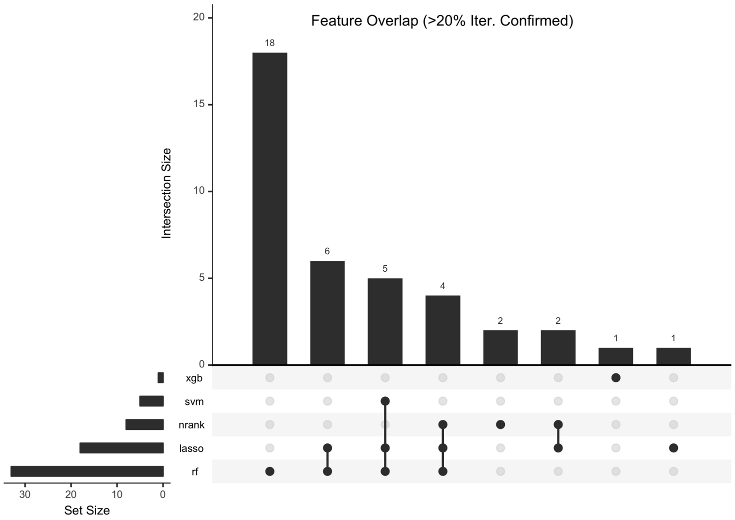


Figure 13: Feature overlap among features confirmed in at least 20% of bootstrap runs, across bootstrap runs with 5 importance metrics

```
                         type.measure = "class", intercept = FALSE)
    #Select lambda min.
    message("selecting lambda min...")
    lambda.min <- cv.fit$lambda.min

    #Fit the full dataset.
    message("fitting whole dataset")
    lassofit <- glmnet(x, y, family = "binomial", standardize = FALSE,
                        lambda = grid, alpha = 1, intercept = FALSE)

    #Extract the coefficients
    lassoimp <- predict(lassofit, type="coefficients", s=lambda.min)
    lassoimp <- lassoimp[2:nrow(lassoimp),1]
    return(lassoimp)
  }
```

#### 3B. Random Forest Importance

Random Forest Importance was defined as the mean decrease in accuracy observed for a gene feature, obtainable using `randomForest::importance()` function.

```
impRF <- function(df, classes, ntrees=100, seed=2019){
    require(randomForest)
    set.seed(seed)
    class <- as.numeric(classes)
    rffit <- randomForest(class ~ ., data = as.matrix(df), ntree = ntrees,proximity = TRUE)
    # rfimp <- as.numeric(getRFIvar(rfmodel=rffit))
    rfimp <- importance(rffit)[,1]
    return(rfimp)
  }
```

#### 3C. XGBoost Importance

XGBoost importance was quantified as either the gain assignments to feature genes, or 0 for excluded genes. These are obtainable using the `xgboost::xgb.importance()` function.

```
impXGB <- function(df, classes, seed=2019){
    require(xgboost)
    set.seed(2019)
    message("fitting xgboost model...")
    xgbfit <- xgboost(data = df, label = classes, max_depth = 2,
                      eta = 1, nthread = 2, nrounds = 2, objective = "binary:logistic")
    message("getting xgboost importances...")
    xgbimp <- xgb.importance(feature_names = colnames(df), model = xgbfit)
    message("reformatting xgb importances...")

    xgbimp.format <- c(rep(0, ncol(df)))
    names(xgbimp.format) <- colnames(df)
    for(f in 1:nrow(xgbimp)){
      xgbimp.format[which(colnames(df)==as.character(xgbimp[f,1]))] <- as.numeric(xgbimp[f,2])
    }
    xgbimp.format <- as.numeric(xgbimp.format)
```

```
    names(xgbimp.format) <- colnames(df)

    return(xgbimp.format)
  }
```

##### 3D. SVM Importance

SVM importance was quantified as the final weight assignments for each gene feature.

```
impSVM <- function(df, classes, seed=2019){
    require(e1071)
    set.seed(2019)
    message("fitting svm model...")
    svmfit <- svm(as.factor(classes)~.,
                  data=df,
                  method="C-classification",
                  kernel="linear")
    svmimp <-  t(svmfit$coefs) %*% svmfit$SV
    message("reformatting importances for output...")
    svmimp.format <- svmimp[1,]
    names(svmimp.format) <- colnames(svmimp)
    return(svmimp.format)
  }
```

##### 3E. Consensus Importance

Consenus importance was defined as normalized median absolute rank, or 'nrank', for each gene feature.

```
# CML for Boruta Implementation
impBorutaCML <- function(x=x, y=y, seed, ranksummary="median",
                         algo.opt=c("lasso","rf","svm","xgb")){
    # impCML
    # Get consensus importance ranks from disparate algorithms.
    # Arguments
    # * df (matrix) : Entire dataset matrix (rows = instances, columns = features)
    # * classes (character): categorizations of instances
    # * algo.opt (list): List of valid algorithms to use for consensus
    # * seed (int): Set the seed for reproducibility
    # * ranksummary (string): Either "score", "median", or "mean", the operation used to calculate cons
    # Returns:
    # * Consensus rank, optionally a standard output table of importances for selected algorithms (if s

    # define the test df if provided or if FALSE
    df.test <- as.matrix(x)
    message("proceeding with df of size ",nrow(df.test))
    classes <- y
    print("getting importances...")
    implist <- list()
    implabellist <- c()

    if("lasso" %in% algo.opt){
      # define the trainindices if none provided or FALSE
```

```
    message("performing lasso...")
    imp.lasso <- impLasso(df=as.matrix(df.test),
                          trainindices=seq(1,nrow(df.test)),
                          classes=classes)
    implist[["lasso"]] <- imp.lasso
    implabellist <- c(implabellist, "lasso")
  }
  if("rf" %in% algo.opt){
    imp.rf <- impRF(df=as.matrix(df.test), classes=classes)
    implist[["rf"]] <- imp.rf
    implabellist <- c(implabellist, "rf")
  }
  if("xgb" %in% algo.opt){
    imp.xgb <- impXGB(df=as.matrix(df.test), classes=classes)
    implist[["xgb"]] <- imp.xgb
    implabellist <- c(implabellist, "xgb")
  }
  if("svm" %in% algo.opt){
    imp.svm <- impSVM(df=as.matrix(df.test), classes=classes)
    implist[["svm"]] <- imp.svm
    implabellist <- c(implabellist, "svm")
  }
  # match importance feature label ordering
  if(length(implist)>1){
    for(i in 1:(length(implist)-1)){
      implist[[i]] <- implist[[i]][order(match(names(implist[[i]]),names(implist[[i+1]])))]
    }
  }

  # get importance ranks and rankvars
  if(ranksummary %in% c("median","mean")){
    impranklist <- list()
    # for each rank, calculate normalized rank position
    message("computing normalized rank importances...")
    for(i in 1:length(implist)){
      algoname <- names(implist)[i]
      impvals <- implist[[i]]
      # naive rank
      nranki <- rank(abs(impvals))
      # adj1: compress rank lower bounds
      min.ranki <- min(nranki)
      nranki1 <- nranki
      nranki1[nranki1==min.ranki] <- min(nranki1[!nranki1==min.ranki])-1
      # adj2: normalize rank scale (dif.min/range)
      nranki2 <- nranki1
      nranki2 <- (nranki2-min(nranki2))/(max(nranki2)-min(nranki2))
      # always compute absolute ranks
      impranklist[[algoname]] <- nranki2
    }
    medrank <- c()
    meanrank <- c()
    # iterate on features
    for(r in 1:ncol(x)){
```

```r
      rvalr <- c() # get feature-level ranks
      for(i in 1:length(impranklist)){
        rvalr <- c(rvalr, impranklist[[i]][r]) # append ranks
      }
      # append summarized feature-level ranks, to return
      medrank <- c(medrank, median(rvalr))
      meanrank <- c(meanrank, mean(rvalr))
    }
    # final importance value is directly proportional to summarized rank
    if(ranksummary=="median"){
      lr <- medrank
    } else{
      lr <- meanrank
    }
    message("completed all tasks! Returning...")
    lr
    return(lr)
  }
  return()
}
```
